# Butyrate ameliorates inflammation in colon biopsy samples of IBD patients and experimental colitis in mice involving RNA binding protein, AUF1-IL-27 axis and accelerating B1a to B10 polarization

**DOI:** 10.1101/2024.05.24.595646

**Authors:** Aaheli Masid, Oishika Das, Diganta Roy, Ankita Dutta, Sohini Sikdar, Atanu Ghosh, Arpan Banerjee, Ujjal Ghosh, Sutanu Acharya Chowdhury, Sankhasubhro Majumder, Mohammad Yahya, Surajit Sinha, Moumita Bhaumik

## Abstract

The pathophysiology of Inflammatory Bowel Disease (IBD) is significantly influenced by the decline in B regulatory (B10) cells, which produce IL-10. Therefore, it is important to identify the key genes and pathways that regulate the B10 cell generation in order to develop more effective therapies. Here, we have shown that one of the short chain fatty acid, butyrate regulates the expression of RNA binding protein, AUF1 which is responsible for increasing the half-life of p28 mRNA, coding for p28 protein which associates with overexpressed EBI3 and forms functional IL-27. This effect is mediated through AUF1 binding to 3’UTR of IL-27p28 mRNA. As a consequence, IL-27 signals splenic CD19^+^CD5^+^ (B1a) cells but not CD19^+^CD23^+^ (B2) cells to polarize to B10 cells. We proved the importance of AUF1 and the sequential downstream players in unique cell penetrating morpholino induced AUF1 knockdown (AUF1-KD) in mice, establishing the roster of events in splenic B1a cells: butyrate-AUF1-IL-27-IL-10. We showed that there was a significant decrease in AUF1, IL-27 and IL-10 expression in the colon biopsy of IBD patients compared to non-IBD control. We have used DSS induced colitis in mice as a surrogate of IBD in human and showed the reduction in AUF1 in spleen and colon could be correlated with the decrease in IL-27 and B10 cells in spleen and mesenteric lymph nodes which were reversed with butyrate treatment. We further established AUF1 as the role player by showing adoptive transfer of butyrate stimulated B1a cells from wild type mice conferring protection against colitis while adoptive transfer of butyrate stimulated B1a cells from AUF1 KD mice failed to suppress the disease. Finally, we propose that butyrate driven B1a cells as a glimmer of new hope of therapeutic possibility against colitis.

## Introduction

Variety of pathobiological pathways epitomize complexities of colitis, but steady barrage of reports resonates that the altered immune homeostasis due to gut resident microbial dysbiosis posits short-chain fatty acids (SCFAs) like butyrate, propionate, and acetate to wrench local inflammatory response at bay (1). There are multiple narratives on human studies stating the beneficial role of butyrate in remission of variety of gut origin symptoms (2), (3). Albeit T-cells have created an endearing hinge, but the landscape of B-cells acceding attention in recent time to understand colitis pathogenesis. The murine model of colitis displays poorly regulated B-cells to cause exacerbated inflammation by blocking T_reg_ function (4, 5). During colitis, number of T_reg_ in gut associated lymphoid tissues in B-cell deficient mice reduced significantly which is restored upon B-cell transfer (6). Exploiting bone morrow chimera system, it was shown that in absence of IL-10 produced by B-cells (B10 cells), mice were unable to show cure from colitis (7). In human, B-cell depletion for unrelated diseases using anti-CD20 (rituximab) reported to either aggravate colitis or caused spontaneous colitis (8). The most studied and significant subpopulation of B regulatory cells that can produce IL-10 to inhibit the immune system are called B10 cells (9). B cells can be divided into several subclasses that differ in <u>ontogeny</u>, <u>homeostasis</u>, and functionality (10). The earliest B cells to form through development are B-1 cells mainly dwell in the peritoneal cavity or umbilical cord blood (11) and also found in the spleen (12). B-1a cells but not B-1b cells display cell-surface CD5 and functions by suppressing endogenous pathogens or microbiota (13), (14). B2 cells, which are characterized by presence of CD23 and the major constituent of B cells, are short-lived recirculating cells and collaborate with helper T cells to generate specific antibodies (15). Maintenance of mature B-cell homeostasis is reported to be dependent on AU-rich element RNA-binding factor 1 (AUF1). AUF1 encoded by *AUF1*/ *hnrnpd* gene and have four isoforms AUF1^p45^, AUF1^p42^, AUF1^p40^ and AUF1^p37^ generated through differential splicing (16). Based on the studies in AUF^-/-^ mice showing follicular-B cells reduced by two-fold whereas marginal zone-B-cells remained unaltered in as compared to wild type counterpart. Different B-cell subsets showed different levels of AUF-1 isoform expression. AUF^-/-^ mice showed unaltered cellular repertoire in lymph node, thymus, bone marrow coupled but two-fold reduction in splenocytes as compared to wild type mice. Interestingly, T-dependent antigen response is comparable whereas T-independent antigen response showed relatively compromised in AUF^-/-^ mice as compared to wild type counterpart (17).

AUF1 binds to AU-rich elements within the mRNA mostly at 3′ untranslated region (3′UTR), leading to either stabilizing mRNAs such as encoding TWIST-1 (18), Jun-D (19), c-myc (20) or destabilising the mRNAs encoding such as TNF-α (21), GM-CSF (22), COX-1 (23), IL-2 (24). Overproduction of TNF-α, in AUF1 knockout mice accounts for higher susceptibility to septicaemia (25). Previously we showed that AUF-1 is increased due to butyrate treatment (26) in mice liver and AUF-1 knock down using a cell penetrating morpholino in mice, inhibits butyrate mediated hypo-cholesterolemic effects in mice (26).

Butyrate activates RNA binding protein, AUF1 (26), is known to regulate cytokine mRNA stability (27), (28), (29). It is recently reported that B-1a cells (30) and Treg produce IL-27 (31). There is a report of IL-27 production from B1a (32) and IL-27 induces IL-10 production from Treg cells (33). The fact that IL-27 as a biologic to suppress auto immune processes (34) offers a valuable framework to address similar question in gut homeostasis. When IL-27 signalling is blocked during bacterial infection, enhanced pro-inflammatory response initiated through downregulation of IL-10 (35). Thus, one is left with a notion that AUF-1 and IL-27 may have a cross talk either directly or indirectly which may govern the colitis pathology. Interestingly there is a report that IL-27 can rescue the disease phenotype of acute severe colitis via IL-10 which is believed to be T-cell independent (34). IL-27 is encoded by two distinct genes, EBI3 and IL-27p28 and their heterodimer forms IL-27 and its receptor consists of heterodimer of IL-27Rɑ (WSX-1) and gp130. IL-27p28 also called IL-27A or IL-30 can function independently without EBI3 (36).

We narrate here, the mechanism by which butyrate stabilizes IL-27 mRNA to polarize B-1a cells to B10 and suppresses DSS induced colitis in mice. We have shown that butyrate modulates AUF1 leading to increased half-life of IL-27p28 mRNA, resulting in the polarization of B-1a cells to B10. To provide further leverage in our understanding, we used morpholino to knock-down AUF1 in mice, and showed that butyrate indeed exploits AUF1 to accomplish its effect on B-1a cells. We have shown in IBD patients and DSS induced colitis in mice that AUF1 and IL-27 expression in colon tissue is significantly decreased. Remarkably, we found that transfer of butyrate stimulated B1a cells from wild type mice conferred protection against DSS induced colitis while adoptive transfer of butyrate stimulated B1a cells from AUF1 knock down mice could not suppress the disease. To our knowledge, we finessed here for the first time that butyrate orchestrates its effect through RNA binging protein AUF-1 to increase IL-27p28 mRNA stability and resulting biologically active IL-27 which polarizes B1a cells to polarize into B10 cells as an effusive entity to control colitis.

## Results

### Butyrate but not propionate or acetate treatment in mouse splenocytes induced IL-10

We endeavoured to learn the ability of acetate, propionate and butyrate (SCFAs) to induce IL-10 production from mouse splenocytes. The splenocytes were exposed to 10 μM and 100 μM of each of the SCFAs in the culture condition and resulting IL-10 production was assayed by ELISA. Our results showed that there was hardly any induction of IL-10 in splenocytes in response to any one of the SCFAs at 24h (Fig 1A) but increased significantly with all three SCFAs at 10 μM and 100 μM at 48 h compared to control (Fig 1B). Interestingly at 72 h, treatment with10 μM acetate and propionate did not show enhancement in IL-10 production but at 100 μM there was low but significant increase in IL-10 induction as compared to control (Fig 1B). Under identical scenario, butyrate treatment caused 1.3 and 1.8-fold higher at 10 and 100 μM of butyrate respectively as compared to the corresponding doses of acetate and propionate (Fig 1C). Therefore, further experiments were conducted with 100 μM at 72 h timepoint.

**Fig. 1:**
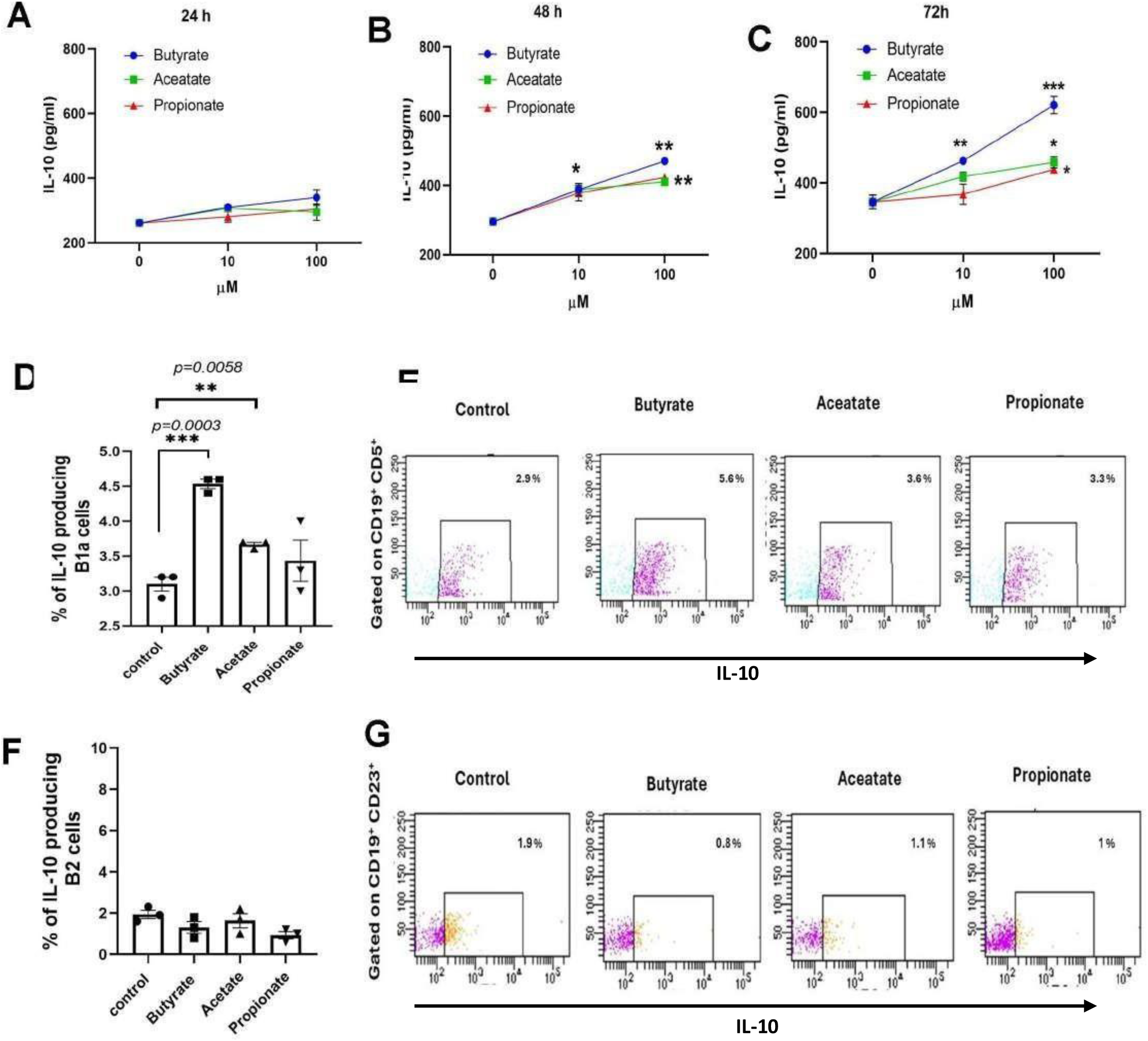
Effect of butyrate or propionate or acetate (SCFA) on mouse splenocyte on IL-10 production and phenotypic characterization of IL-10 producing cells. Splenocytes were treated with either butyrate or propionate or acetate at a concentration of 10 uM and 100 µM, where without SCFAs were used as control and resulting IL-10 production in the culture supernatant was measured by ELISA at 24h (A), 48h (B) and 72h (C). Having established that 100 uM of SCFAs caused significantly higher IL-10 induction in splenocytes compared to without SCFAs, rest of the studies were carried out with 100 µM. For phenotypic characterization of IL-10 producing cells, splenocytes were treated with 100 µM of each SCFAs for 72 h where monensin treatment was added during the last 8 h of culture. B1a cells and B2-cells in the splenocytes were characterized as CD19^+^CD5^+^ and CD19^+^CD23^+^ respectively and on the gated population intracellular IL*-*10 production was assayed stanning with anti-IL-10 mAb and results were expressed % IL-10 producing cells in B1a (D and E) and B2 cells (F and G). Gating strategy is shown in supplementary Fig S1. All experimented were repeated thrice (n=3) and represented as Mean ± SE. *p < 0.05; **p < 0.01; ***p < 0.001 with respect to control calculated by student’s t-test.

### B1a-cell but not B2 cells as a source of IL-10

We plan to navigate into the orbit of B-cell based on the fact that a unique phenotype of IL-10 producing B-cell posits as regulatory B-cells. To shed light on the process, two sub-lineages of B-cells: B1a cells as CD19^+^CD5^+^ and B2 cells as CD19^+^CD23^+^ (37) were considered. Splenocytes were cultured without and with 100 μM butyrate for 72 h and B1a and B2 cells were gated and percentage of IL-10 producing cells was enumerated by intracellular cytokine assay. Our results showed that IL-10 producing B1a cells increased by 1.5-fold due to butyrate treatment whereas acetate but not propionate showed low but significant increase in frequencies (Fig 1D, E). Interestingly, B2 cells did not show any such increase in response to treatment with any one of the above SCFAs (Fig 1 F, G). Henceforth gated population of IL-10 producing B1a cells are defined as B1a cells for convenience. Gleaned from above studies accedes a quest to cultivate link between butyrate and B1a cells whether it is a direct effect or indirect.

### rIL-27 increases the frequencies of IL-10 producing B-1a cells but not B2 cells

To address the link, if any, splenocytes were treated with 20 ng/ml rIL-27 as reported earlier (38), (39) and after 72h, frequencies of B1a cells but not B2 cells was enumerated. It was observed that IL-10 producing B1a cells but not B2 cells increased by 3-fold with respect to control (Fig S2 A, B). Based on this information we enquired if butyrate has any effect on IL-27 expression. Our results showed that there was hardly any change of EBI3 expression in splenocytes in response to 10μM and 100μM butyrate treatment but IL-27p28 expression showed dose dependent significant increase compared to control (Fig 2A). The increase in IL-27p28 were further verified by ELISA and western blot. The estimation in the increase of IL-27p28 protein with butyrate treatment by ELISA and western blot analysis gave similar results (Fig 2B, C, D).

**Figure 2:**
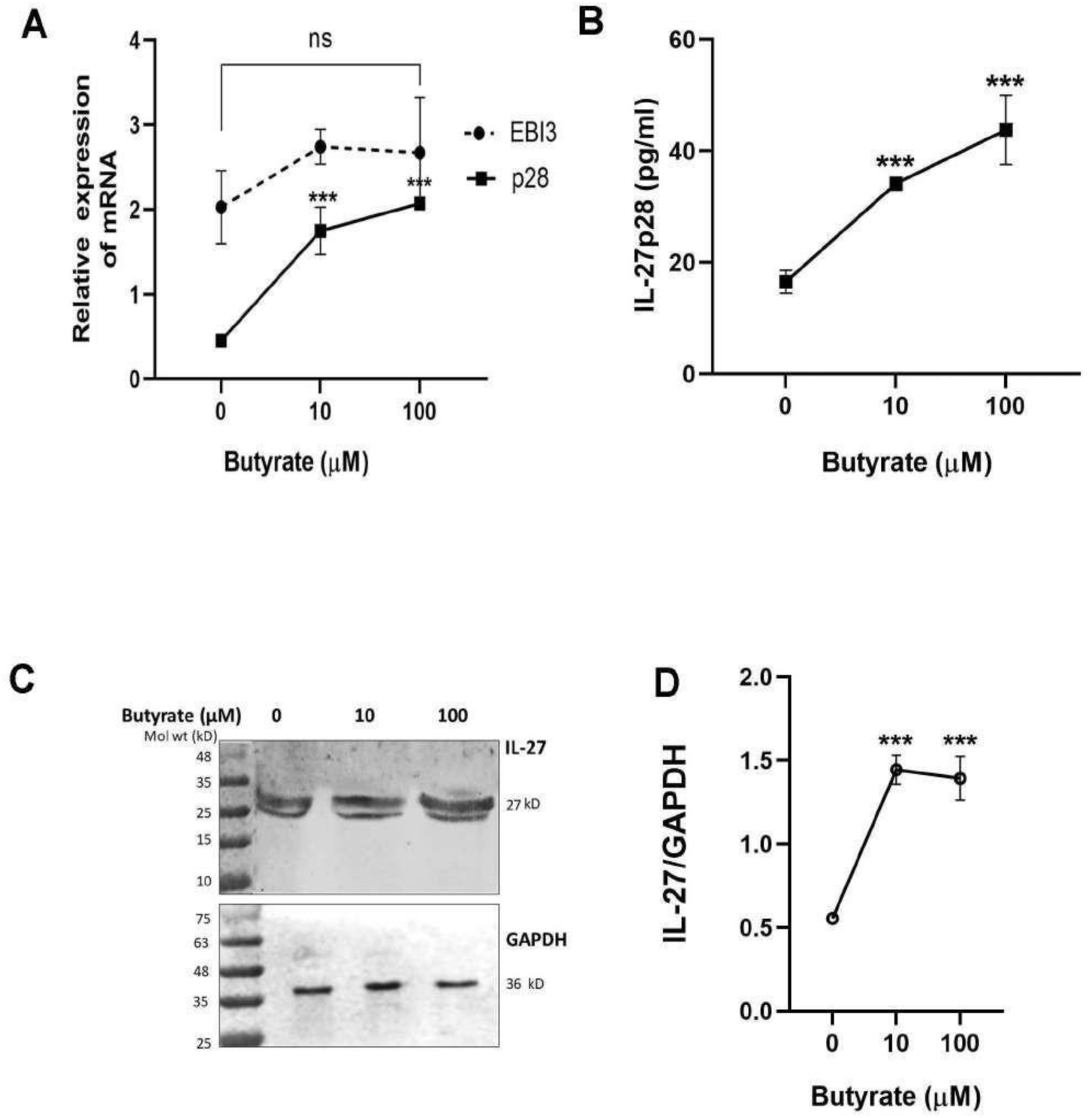
Effect of butyrate on IL-27 production by normal mouse splenocytes. Splenocytes were treated with butyrate at a concentration of 10 µM and 100 µM for 72 h. The expression of p28 and EBI3, the two subunits of IL-27 was determined by qPCR (A) and corresponding IL-27p28 was determined in the culture supernatant (B), expression of IL-27 by western blot (C) and corresponding densitometry of the western blot using ImageJ showing relative expression of IL-27 with respect to GAPDH (D). All experimented were repeated thrice (n=3) and represented as Mean±SE. p values calculated with respect to control by student’s t-test.

With an objectivity to fit into the triptych of the following: butyrate, IL-27 and B1a cells, the following array of experiments were considered.

#### Both butyrate or IL-27 increased B10 cells in similar magnitude

To find the revelatory relation, if any, between butyrate and IL-27, the magnitude of B10 cells induction in splenocytes in response to either butyrate or rIL-27 treatment was evaluated as functional readout. It was observed that butyrate treatment caused 2.4-fold increase in B10 cells which was reduced to basal level in the presence of anti-IL-27 mAb. Similarly, treatment of splenocytes with rIL-27 alone also rendered generation of B10 cells essentially similar magnitude as observed with butyrate (Fig 3A). This observation harnessed the interiority of butyrate effect on B1a cells was IL-27 dependent. The higher expression of CD1d in B-cells is considered as a marker of B_reg_ cells (40) and to assess the regulatory nature of B1a cells, expression of CD1d in IL-10 producing B1a cells were studied. It was observed that IL-10^+^CD1d^+^B1a cells frequency was increased significantly in the presence of rIL27. Furthermore, butyrate also increased the frequencies in similar magnitude as with rIL-27 which was reduced to the basal level in the presence of anti-IL27 Ab and corresponding flow cytometric analysis (Fig 3B, C).

**Figure 3:**
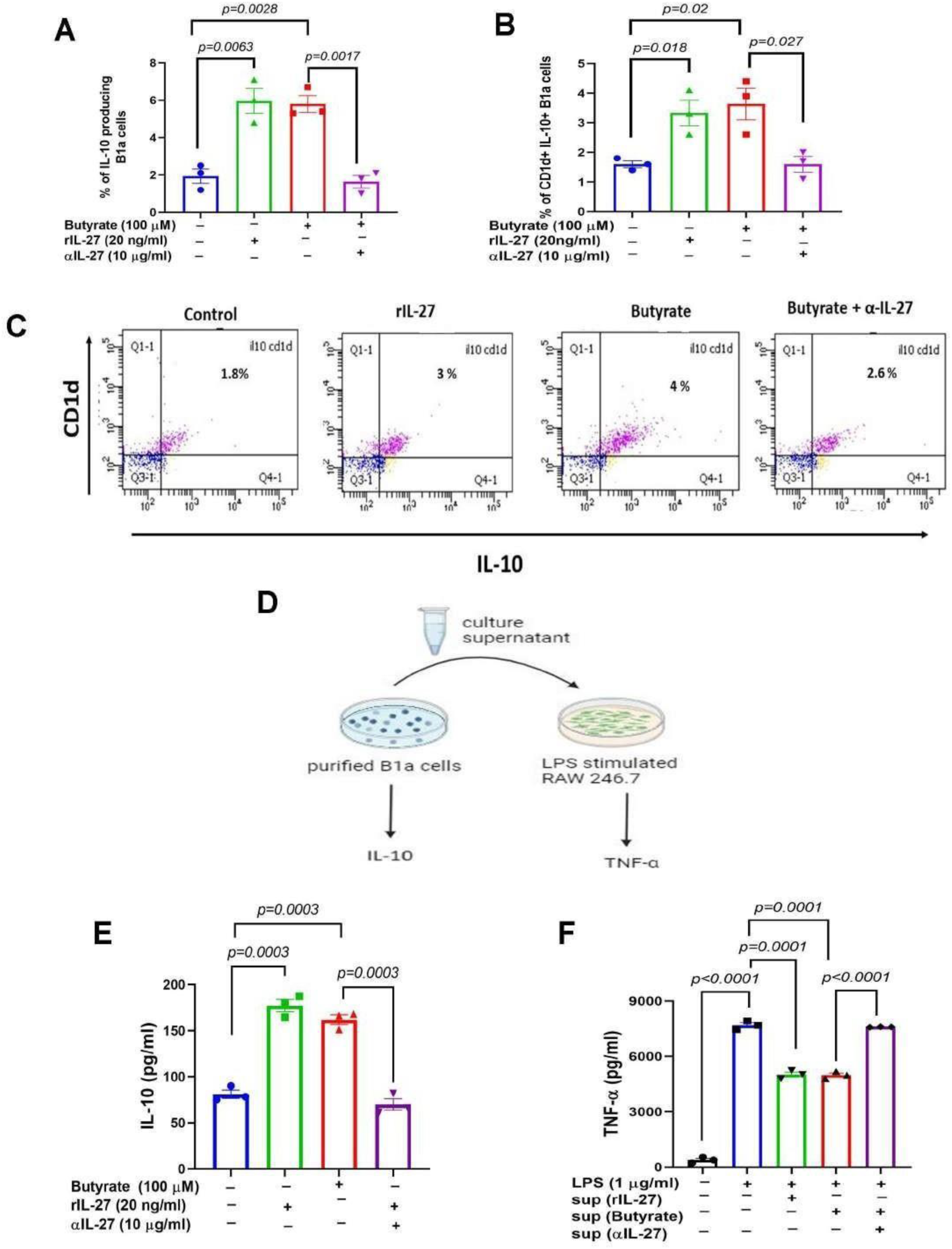
Butyrate or rIL-27 induced IL-10 production from splenocytes and B1a cells and functional assay of supernatant of B1a cells as inhibitor of TNF-α production. Splenocytes treated either without (**□)** or with butyrate (100 µM) (**□**) or rIL-27 (20 ng/ml) (**□**) or butyrate (100 µM) plus α-IL-27 mAb (10 µg/ml) **(□**) for 72 h and monensin was added to last 8 h of treatment. Thereafter cells were stained with antibodies against CD19, CD5 and CD1d. The percentage of IL-10 positive B1a cells in the splenocytes were detected after staining with anti-IL-10 Ab and gated on CD19^+^CD5^+^ cells (A). The percentage of CD1d^+^IL-10^+^ B1a cells was detected by gating CD19^+^CD5^+^ cells (B). The representative dot plots showing CD1d^+^IL-10^+^ B1a cells gated on CD19^+^CD5^+^ cells (C). The CD19^+^CD5^+^ cells were purified from untreated splenocytes and were cultured *in vitro* without (**□**) with either butyrate (100 µM) (**□**) or rIL-27 (20 ng/ml) (**□**) or butyrate (100 µM) plus α-IL-27 mAb (10 µg/ml) (**□**) for 72 h. A part of the supernatant was collected to measure IL-10 production by ELISA and the other part was added to 24 h LPS (10 ng/ml) stimulated RAW264.7 cells which were further cultured for 24 h. The TNF-α was measured by ELISA from the resulting supernatant from LPS stimulated RAW264.7 after 24 h. The diagram represents experimental design (D). The ELISA results of IL-10 (E) and TNF-α (F) are presented. The pre and post sorting is represented in supplementary figure S3. All experimented were repeated thrice (n=3) and represented as Mean±SE. p values calculated with respect to control by student’s t-test.

#### Purified B1a cells produce IL-10 in response to butyrate or IL-27

To show that butyrate or rIL-27 activates B1a population directly to produce IL-10, B-1a cells (CD19^+^ CD5^+^) were purified from whole splenocytes and cultured *in vitro* for 72 h either with rIL-27 or butyrate or butyrate + anti-IL-27 mAb and resulting IL-10 production was measured in culture supernatant by ELISA (Fig 3D). It was observed that there was a significant increase in IL-10 in culture supernatant due to rIL-27 or butyrate treatment as compared to untreated control. As expected, butyrate treatment in the presence of anti-IL-27 mAb, IL-10 production returned to basal value (Fig 3E). To establish credible evidence that B1a cells renders anti-inflammatory property upon butyrate treatment, functional assay was envisaged considering TNF-α as an inflammatory cytokine.

#### Butyrate or rIL-27 treated B1a cell supernatant inhibits TNF-α production

Functional analysis was done to study the ability of IL-10 produced by B-1a cells having anti-inflammatory property or not, the supernatant of cultured B1a cells treated either with butyrate or rIL-27 or butyrate + anti-IL-27 mAb and measured LPS stimulated RAW 246.7 cells derived TNF-α production. The LPS stimulation of RAW 246.7 cells caused 15-fold increase in TNF-a production compared to control. Addition of culture supernatant of either from rIL-27 or butyrate treated B1a cells caused significant reduction in TNF-α production but culture supernatant from butyrate treated B1a cells generated in the presence of anti-IL27 reversed the inhibitory effect (Fig 3F).

The above experiments deftly mirroring as follows: butyrate stimulates B1a cells to produce IL-27 which in paracrine manner producing IL-10 and this resulting IL-10 inhibited TNF-α production from LPS stimulated RAW264 cells. Now the tale unfolds with a notion of reciprocity between IL-27 and TNF-α and butyrate may deftly stroke such a reciprocity.

### Butyrate reciprocally regulates IL-27p28 and TNF-α mRNA stability

We studied the decay of IL-27p28 and TNF-α mRNA to shed light on their reciprocity. The decay of mRNA was measured in terms of their half-life (t_1/2_) in minute which was measured after blocking *de novo* RNA synthesis with Actinomycin D in splenocytes with and without butyrate treatment. It was observed that t_1/2_ of IL-27p28 RNA with and without butyrate treatment were 53.75± 0.7 m and 118 ± 21.5 m respectively (Fig 4A) and corresponding t_1/2_ for TNF-α mRNA was 52.11± 1.26 and 23.1± 2.23 respectively (Fig 4B). Thus, a reciprocity was noted as regard to t_1/2_ of IL-27p28 mRNA and TNF-α mRNA in splenocytes in response to butyrate. This study showed that butyrate treatment caused dual events in tandem: increased and decreased stability of IL-27p28 RNA and TNF-α mRNA respectively. As a control, t_1/2_ of housekeeping gene, GAPDH mRNA regardless of butyrate treatment remained unaltered (Fig 4C). Since, butyrate activates RNA binding protein, AUF1 (26), that exhibits characteristics of a trans-acting factor contributing to the stability and decay of many cellular mRNAs and therefore it was of automatic choice to study linking IL-27 with AUF-1.

**Fig 4:**
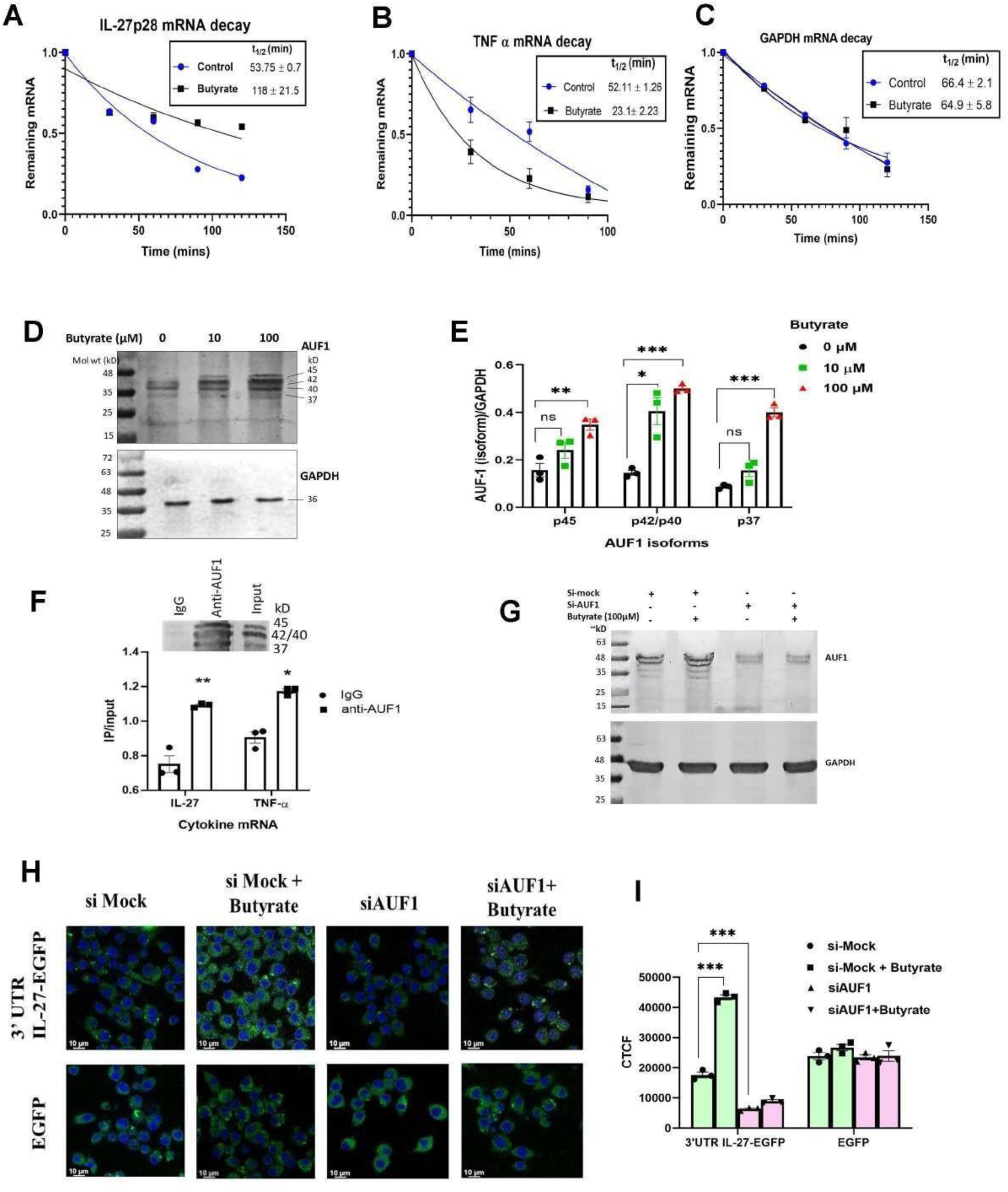
Butyrate regulates p28 mRNA stability by binding to 3’UTR of IL-27p28 mRNA. Splenocytes treated with or without 100 µM butyrate for 72 h. Then Actinomycin D (10 μg/ml) was added to the splenocytes and RNA was extracted at indicated timepoints to measure mRNAs by qPCR. The remaining IL-27p28 mRNA levels (A), TNF-α mRNA (B) and GAPDH mRNA (C) were measured by qRT-PCR and normalized to the results at 0 min and represented as a function of time. AUF1 isoform expression in the splenocytes after 72 h of treatment with 10 µM and 100uM butyrate by western blot (D). The corresponding densitometry showing relative AUF-1 isoform expression with respect to GAPDH (E). Whole cell lysates extracted from splenocytes were used for RNA-IP with anti-AUF1 Ab (inset, F). Both p28 and TNF-α mRNA were detected by qPCR in RNA extracted from the RNA-IP complex (F). RAW246.7 cells were transfected with either si-AUF1 or mock RNA and further treated with or without butyrate for 72 h. The AUF1 expression was measured by western blot (G). EGFP constructs carrying the 3′UTR of gene encoding p28 (3′UTRp28) or empty vector (pEGFP) was transfected into siRNA transfected cells and EGFP fluorescence was measured by fluorescence microscopy in si-mock, si-mock+ butyrate, si-AUF-1 and si-AUF1+butyrate (H). The relative fluorescence intensity was determined (I). All experimented were repeated thrice (n=3) and represented as Mean±SE. p values calculated with respect to control by student’s t-test.

### Butyrate upregulates AUF1 and interaction of AUF-1 with TNF-α and IL-27p28 mRNA

To show the effect of butyrate, if any, on AUF-1, splenocytes treated with or without butyrate and the expression of AUF-1 isoforms (p45, p42, p40 and p37) were studied by western blot (Fig 4D). The western blot showed that the AUF1 isoforms: p45, p42, p40 and p37 were increased upon butyrate treatment (Fig4 D). It was clear from densitometry analysis that at 10 µM butyrate treatment p42/p40 and p45 showed significant increase compared to control. However, 100 µM butyrate treatment, all the AUF1 isoforms (p45, p42/p40 and p37) were increased 2-fold or more compared to control (Fig 4 E). Now to reveal if IL-27p28 mRNA remained associated with AUF1, we performed RNA immunoprecipitation (RNA-IP) with anti-AUF1 antibody in whole splenocytes without any treatment. The western blot showed the presence of AUF1 in anti-AUF1 pull down samples but not in control IgG pull down samples (Fig 4F, Inset) and qPCR analysis further showed that both IL-27p28 and TNF-α mRNA are present in anti-AUF1 pull down and not in IgG pulled down samples (Fig 4F). Since our study is to evaluate the mechanism of butyrate mediated IL-10 induction exploiting IL-27p28 mRNA stabilization we sought to understand where does AUF1 binds to IL-27p28 mRNA.

### AUF-1 binds to 3’-UTR of IL-27p28 mRNA

As AUF1 mostly binds to the 3’UTR of its target mRNAs (41), (18) we seek to understand if it also binds to the 3’UTR of IL-27p28 mRNA. Therefore, we cloned the IL-27p28 3′UTR immediately upstream of EGFP gene and transfected it into RAW246.7 cells. To show that AUF1 stabilizes IL- 27p28 mRNA by binding to the 3’-UTR, a co-transfection experiment was performed. The cells were transfected either with pEGFP-3’UTRp28 or pEGFP and the resulting EGFP fluorescence was analysed by fluorescence microscopy. Following the second round of transfection either with si-mock or siAUF1, the cells were treated with or without butyrate generating four experimental sets from left to right as follows: (i) si-mock, (ii) si-mock plus butyrate, (iii) siAUF1 and (iv) siAUF1 plus butyrate. The western blot showed the effective knock down of AUF1 expression which was not reversed even after butyrate treatment (Fig 4G). To bring further credence to our observation, transfection in RAW246.7 cells was done with pEGFP-3’UTRp28 (Fig 4H, upper panel) or pEGFP (Fig 4H, lower panel). In the upper first panel from left only si-mock showed basal level of fluorescence expression which was increased significantly with si-mock plus butyrate treatment (second panel). In the third and fourth panel either siAUF-1 or siAUF-1 plus butyrate treatment showed significant decrease in fluorescence as compared to si-mock treatment. The fluorescence intensity was quantified and expressed as corrected total cell fluorescence (CTCF). The fluorescence intensity of 3’ UTR IL27 EGFP and EGFP as per above specified conditions was expressed in bar diagram (Fig 4I).

### Relevance of AUF-1 knock down on p28mRNA expression, stability, translational regulation and IL-10 producing B1a cells

To bring further strength to our notion on the importance of AUF-1 in the butyrate mediated effects, we opted to generate AUF1 knock down (AUF1-KD) in mice using novel cell penetrating morpholino oligomer of GMO-PMO (MO), to knock down AUF-1 in mice for further phenotypic analysis.

#### GMO-PMO driven knock down AUF1 isoforms in mice

To bring further strength to our notion on the importance of AUF-1 in the butyrate mediated effects, we opted to generate AUF1 knock down (AUF1-KD) in mice using novel cell penetrating morpholino oligomer of GMO-PMO. Here, anti-sense morpholino oligomer, GMO-PMO specific to AUF1 is denoted as AUF1-MO and corresponding scramble is denoted as scramble-MO (S-MO). The expression of AUF1 in the spleen cells from all the groups: control, S-MO, S-MO + butyrate, AUF1-MO and AUF1-MO+butyrate was analysed by western blot (Fig 5A, B). As expected, there was no difference in AUF-1 expression in the splenocytes of control and S-MO mice. Likewise, unsurprisingly S-MO + butyrate treatment caused significantly higher expression of all the isoforms of AUF-1 compared to S- MO. On the other hand, AUF-1 isoforms expression was hardly detectable in AUF-1-MO mice which was not restored upon butyrate treatment. (Fig 5B). Therefore, it was important to explore the impact of AUF1-Knock down (AUF-1KD) on IL-27p28 expression and its stability in in vivo system.

**Figure 5:**
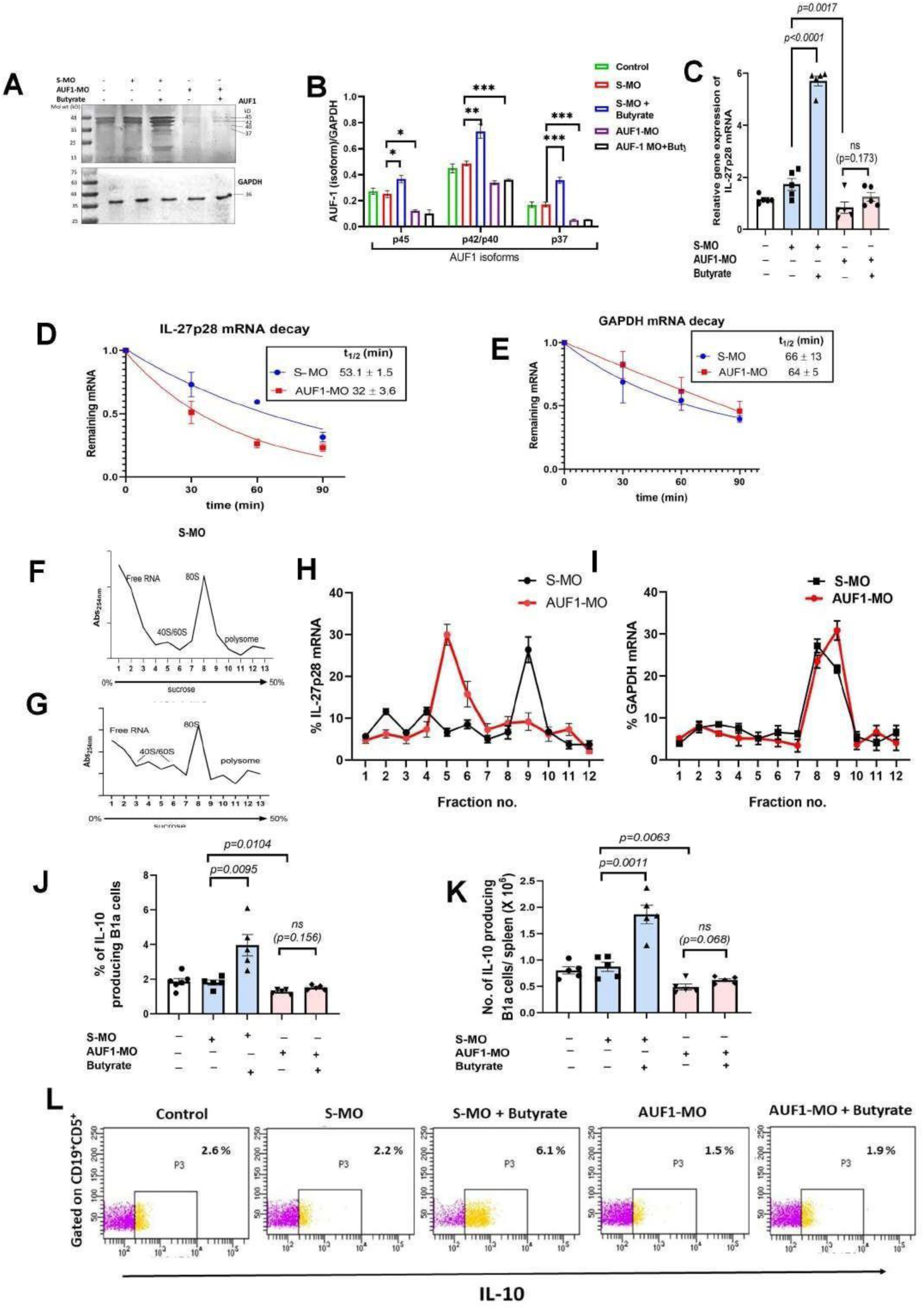
Effect of AUF1 knockdown (AUF1-KD) by Morpholino oligomer (GMO-PMO) on the half-life of IL-27p28 and B10 polarization. Mice were divided randomly into five groups having n=5mice/group: (i) mice without any treatment (ii) mice receiving scramble-MO (S-MO-mice), (iii) mice receiving S-MO fed with butyrate supplement (butyrate-S-MO-mice), (iv) mice receiving AUF-1-MO (AUF-1-MO-mice) and (v) receiving AUF-1- MO fed with butyrate supplement (butyrate-AUF-1-MO-mice). Western blot analysis of AUF1 in spleen in groups i-v (A), corresponding densitometric analysis of AUF1 isoforms expression normalized to GAPDH (Data represent >3 independent experiments. (B). Expression of IL-27p28 mRNA in spleen determined by qPCR in groups i-v (C). Splenocytes isolated from AUF1-MO mice and S-MO mice were treated with or without 100 µM butyrate for 72 h and Actinomycin D (10 μg/ml) was added. Thereafter RNA was extracted at indicated timepoints to study mRNA expression. The remaining p28 mRNA levels and GAPDH mRNA were measured by qPCR and normalized to the results at 0 min after Actinomycin D treatment (D, E) (n=3, t-test). To quantify the mRNA associated with ribosomal fractions from splenocytes were analysed by 0–50% continuous sucrose density gradient fractionation. Ribosomal RNA content, measured at 254nm, from S-MO mice (F) and AUF1-MO mice (G) were plotted against fraction numbers. RNA isolated from selected fractions were analysed by qPCR using appropriate primer sets for IL-27p28 (H) and GAPDH (I) (n=3). The splenocytes isolated from each mouse were treated with monensin for 8 h. Thereafter the cells were stained with antibodies against CD19, CD5 and IL-10. The intracellular IL-10 positive cells were detected with their respective antibodies, gated on CD19^+^CD5^+^ cells(L). Gating is shown in supplementary figure S4. The frequency (J) and absolute number (K) of IL-10 producing CD19^+^CD5^+^ cells are represented in the groups i-v. All experimented were repeated thrice with n=5 mice/group. (Data from J and K representation of 3 independent experiments having n=5mice/group, p value one-way ANOVA with post hoc Tukey’s test). *P < 0.05; **P < 0.01; ***P < 0.001; one-way ANOVA with post hoc Tukey’s test.)

#### IL27p28 expression in AUF-1-KD mice

The IL27p28 expression in the splenocytes of mice was studied by qPCR after receiving the following treatment: (i) control (without any treatment), (ii) S-MO, (iii) S-MO + butyrate, (iv) AUF1-MO and (v) AUF1-MO+butyrate. The basal level of IL-27p28 expression remained essentially similar between untreated and S-MO treated groups. As expected, butyrate treatment caused 3-fold increase of IL27p28 expression compared to S-MO mice. Interestingly AUF-1 MO treatment caused significant decrease in IL-27p28 expression as compared to S- MO treated group which was not restored upon butyrate treatment (Fig 5C). The above observation calls for further enquiry of IL-27p28 mRNA half-life (t_1/2_).

#### Measurement of t_1/2_ IL-27p28 mRNA in S-MO and AUF-1KD mice

The process of mRNA degradation of IL-27p28mRNA was directly assessed by measuring the t_1/2_ of IL-27p28mRNA following the addition of Actinomycin D in splenocytes culture isolated from S-MO and AUF1-MO mice. The t_1/2_ of IL-27p28 in spleen cells of in S-MO and AUF1-MO mice was 53± 1.5 and 32 ± 3.6 respectively (Fig 5D). The t_1/2_ of GAPDH mRNA remained essentially unaltered regardless of AUF-1 knock down (Fig 5E). To understand if the decrease in t_1/2_ of IL-27p28mRNA in AUF1-MO mice impacts its translation repression, we performed polysome analysis.

#### Polysome profiling and analysis of translational status of p28mRNA

The role of translational regulation of gene expression, polysome profiling has been exploited in this investigation to infer the translational status of IL-27p28mRNA. The absorbance at 254 nm for the fractions obtained from S-MO and AUF1-MO are showed the location of polysome, monosomes ((80S) or 40S and 60S) and free RNA (Fig 5 F, G). Our study showed that the distribution of IL-27p28 mRNA shifted towards upper fractions containing monosome (80S) or 40S and 60S in AUF1-MO mice whereas in S-MO mice its distribution restricted to polysome (Fig 5 H). As a control we showed that distribution of GAPDH mRNA remained in the polysome site regardless of any form of MO treatment (Fig 5I) showing that AUF-1 impacted translational regulation of IL-27p28mRNA.

#### AUF1-KD mice failed to produce butyrate driven B10 polarization of B1a cells

Since AUF1-KD mice display translational repression of IL-27p28mRNA, we studied its impact in polarization of B1a cells to B10 in terms of IL-10 production on the gated Cd19^+^CD5^+^ population (B1a cells). The results were expressed as percentage of IL-10 producing B1a cells (Fig 5J) and also in absolute number per 10^6^ splenic cells (Fig 5K). As expected, in the S-MO mice receiving butyrate, there was a significant increase in the percentage and total number of splenic B10 cells. In AUF1-KD mice frequency of such cells became comparable to S-MO-group and remained unaltered even after butyrate treatment. The representative dot plots are provided in Figure 5L. It was important to note that the percentage of B1a cells (Cd19^+^CD5^+^) did not change upon butyrate treatment or AUF1 knock down compared to control (Fig supplementary S4). Thereafter we endeavoured to study the frequencies of IL- 10 producing B1a cells in AUF-1-KD mice in response to butyrate or IL-27 treatment.

#### rIL-27 treatment but not butyrate rescue IL-10 production from B1a cells in AUF-1 KD mice

Splenic B-1a cells were sorted and purified from either S-MO or AUF1-KD mice. The purified population of CD19^+^CD5^+^ cells from S-MO mice and AUF1-KD mice having 93.7% and 91.1% purity respectively shown in the dot plots (Fig 6A, B). The cells from the above groups were treated with either butyrate (100 µM) or rIL-27 (20 ng/ml) for 72h and thereafter IL-10 measured in the resulting supernatant. As expected, S-MO-mice showed basal level of IL-10 production which was enhanced significantly upon rIL-27 treatment or butyrate treatment. B-1a cells from AUF1-MO mice showed nearly 2-fold decrease in IL-10 production compared to the B-1a cells of S-MO mice. Further, treatment with butyrate to the B-1a cells from S-MO mice significantly increased IL-10 production but the cells from AUF1-MO mice with butyrate treatment essentially showed no change. Surprisingly that B1a cells from AUF-1-KD mice upon rIL-27 treatment caused copious IL-10 production which was comparable to the IL-10 produced by butyrate or rIL-27 treated B1a cells from S-MO. (Fig 6C). This dissonance in IL-10 production in AUF1-KD mice between butyrate and IL-27 treatment deftly captures a new revelation of AUF-1 independent effect of IL-27 in IL-10 induction in B1a cells.

**Fig 6:**
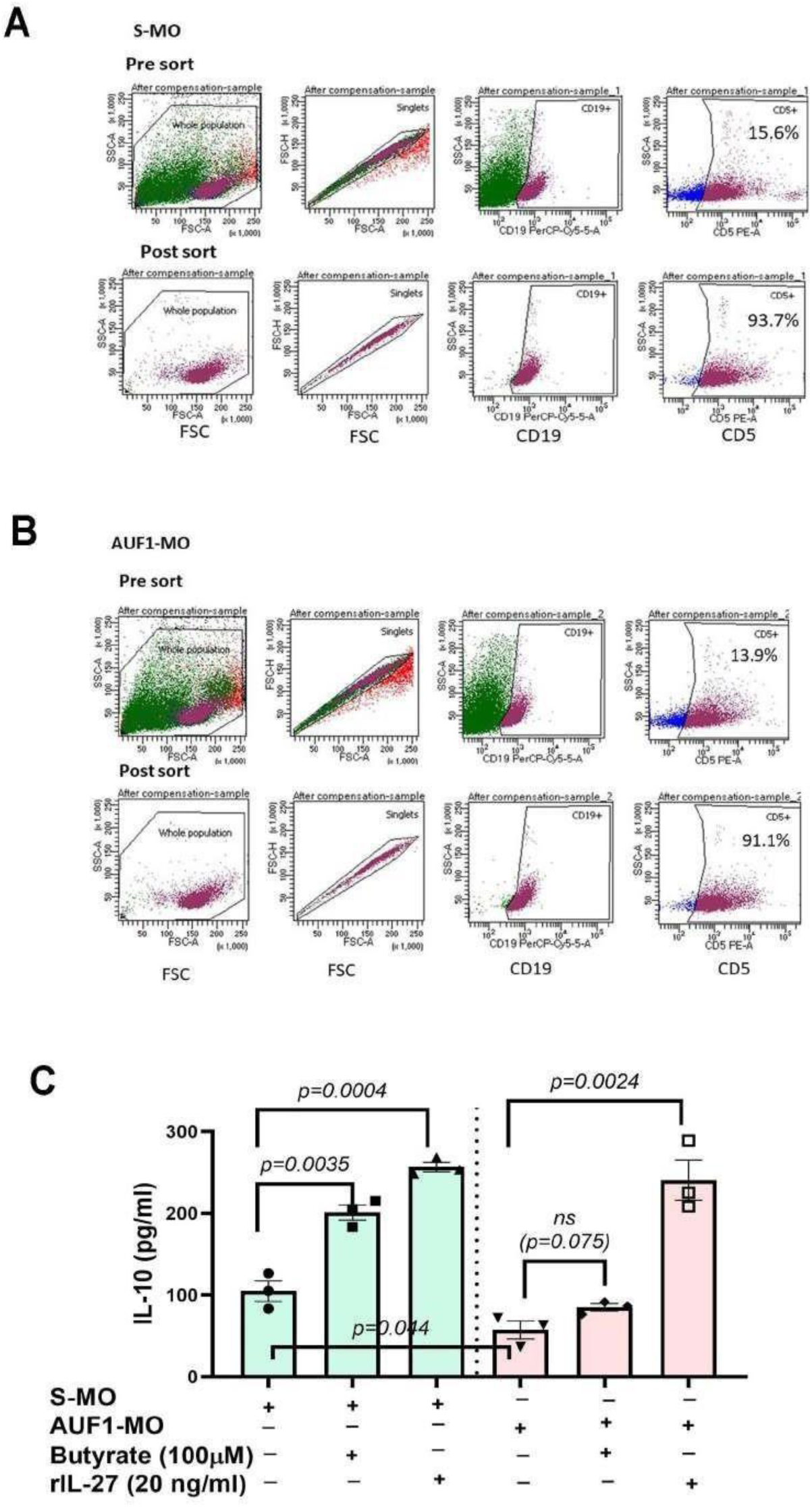
rIL-27 treatment but not butyrate can stimulate B-1a cells from AUF1 knock down mice. The splenocytes isolated from S-MO and AUF1-MO mice were stained with antibodies against CD19 and CD5. The CD19^+^CD5^+^ cells were gated and sorted in FACS. The purity of the sorted CD19^+^CD5^+^ cells from S-MO mice (A) and AUF1-MO mice (B) was determined by pre and post sort flowcytometry. The cells were then treated either with rIL-27 (20 ng/ml) or butyrate (100 µM) for 72 h. Thereafter the resulting supernatant was collected and IL-10 was measured by ELISA (C). Experiment was repeated thrice each time with the cells from 3 independent mice from each group and represented as Mean ± SE. p value one-way ANOVA with post hoc Tukey’s test.

### Analysis of web-source of micro-array data base of AUF-1 knock down human cell line: molecular landscape of inflammatory genes

Evolving landscape from our preceding studies illuminated that AUF-1 KD using AUF-1-MO caused reduction in butyrate driven B10 polarization of B1a cells as compared S-MO treated mice. This observation impelled us to further explore general inflammatory response due to AUF-1 knock down. We searched in the web-source if such information is available in the public domain. The only available AUF-1 knock down cell line to our knowledge is of skin origin of human (<u>GSE138621</u>). Data curated linked with inflammatory processes were derived from the microarray datasets comparing AUF1 knockdown WS1 skin cells vs wild type control. KEGG pathway analysis revealed AUF1-affected networks involved multiple biological processes, including upregulation of IBD (underlined in red Fig 7A). It was observed that multiple mRNAs up/downregulated due to AUF1 knock down and many of them were associated cell adhesion, pathways, Th1/Th2 response, mucosal IgA production and the TGF-β signalling pathway garnering embodiments of gut inflammatory processes (Fig 7A, B).

**Fig 7.**
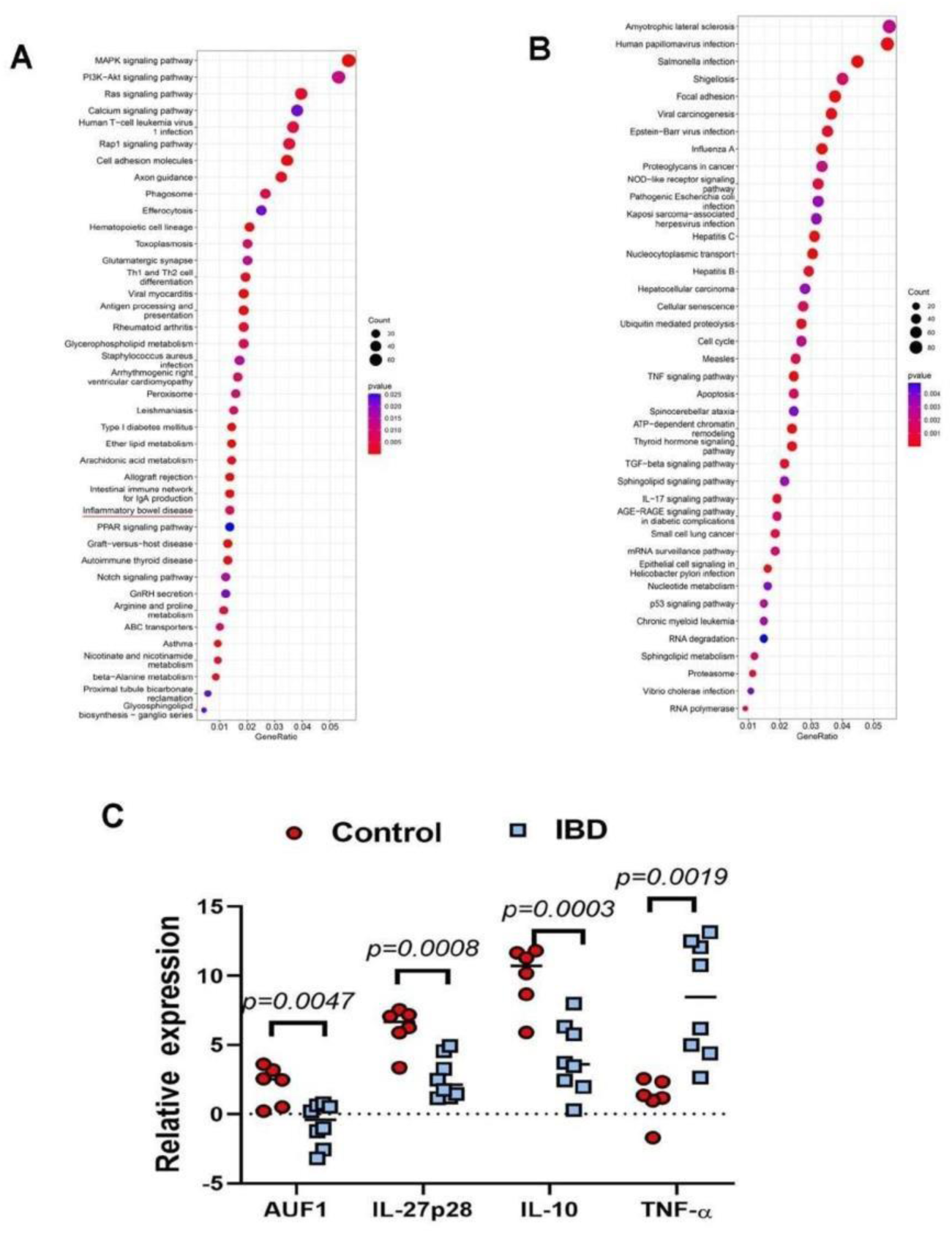
Analysis of web-source of micro-array data of AUF-1 knock out human cell line and AUF- 1 status in Inflammatory Bowel Disease (IBD) patients. KEGG (Kyoto Encyclopedia of Genes and Genomes) pathway analysis of in AUF1 knock out skin cell line, WS1 vs control (<u>GSE138621</u>) was performed showing the upregulated (A) and downregulated pathways (B). The range of colour from blue to red represents the p value as indicated. To gain more insight in real clinical scenario, status of AUF-1 and inflammatory and anti-inflammatory cytokines were studied in the biopsy samples from 6 IBD patients (6 UC, 0 CD) and 6 non-IBD patients were collected from IBD clinic of SSKM hospital. The mRNA expression of AUF1, IL-27p28, IL-10 and TNF-a were determined by qPCR (C). The details of the patients are provided in Table1. Results are representative of triplicate measurements. The data is represented with Median. p values calculated with respect to control by student’s t-test.

### Expression of AUF-1, IL-27, IL-10 and TNF-α in clinical human samples

Early work from patients and IBD mouse models suggested that there is a blurring Th1/Th2 polarization coupled with altered cytokine repertoire. Enriched from the data derived from skin cell line showing signature of inflammatory processes. We analysed AUF1, IL-27, IL-10 and TNF-α expression by qPCR from colonic epithelial biopsies from 10 IBD patients suffering from ulcerative colitis. As controls, we utilized biopsies from 6 otherwise IBD-free individuals who had colonoscopy and whose diagnosis for intestinal diseases were ruled out. The details of the patients are presented in table 1. We observed that the median value of relative AUF1 expression in IBD cohort was nearly fivefold less than the median of control cohort (Fig 7B). Corroborating previous studies, our investigation broadly revealed a 1.89-fold decreased IL-27 expression in IBD patients as compared to the control cohort (Fig 7C). As expected, the IBD patients showed 2-fold decrease in IL-10 production with concurrent fourfold increase in TNF-α compared to control (Fig 7C).

**Table 1:** Details of the patients from whom the colon biopsy samples were collected.

| Patient ID | Age (y) | Sex | UCEIS score | Colonoscopy finding | Samples taken as control or IBD |
| --- | --- | --- | --- | --- | --- |
| 7 | 65 | Male | NA | Normal mucosa, Ileal ulcer | Control |
| 10 | 59 | Male | NA | Normal mucosa | Control |
| 5 | 44 | Female | 5 | Both rectum and sigmoid colon is found with complete loss of vascular pattern with spots and tears. | Active IBD (UC) |
| 6 | 50 | Female | 5 | Complete loss of vascular pattern, large superficial ulcer of 75mm seen | Active IBD (UC) |
| 8 | 24 | Male | -- | Complete loss of vascular pattern seen with multiple superficial ulcers of varying size> 5mm in size. | Active IBD (UC) |
| 12 | 33 | Male | 7 | Mucosa is involved in form of continuous loss of vascular pattern with spots and steaks of blood ahead of scope which can be washed away with jet of water. Multiple superficial and deep ulcer. | Active IBD (UC) |
| 11 | 49 | Female | NA | Normal mucosa | Control |
| 13 |  |  | NA | Normal mucosa, ileal ulcer | Control |
| 15 | 45 | Female | 5 | Complete loss of vascular pattern, expanded lamina propria with inflammatory cells. | Active IBD (UC) |
| 1 | 52 | Male | NA | Normal mucosa | Control |
| 2 | 36 | Female | 7 | ulceration of epithelium, Cryptitis, crypt abscess, mucodepletion. Focal crypt distortion, crypt branching and crypt loss. Lamina propria is expanded with dense amount of inflammatory cells. | Active IBD (UC) |
| 3 | 30 | Male | NA | Chronic sigmoiditis | Control |
| 4 | 58 | Male | 7 | Ulceration in the lining of epithelium, presence of inflammatory cells in lamina propria; no granuloma was found. | Active IBD (UC) |
| 5 | 32 | Male | NA | Normal mucosa | Control |

To ensure that the observed results were not specific to just the human diseases and their matched mouse model of colitis induced by DSS can be used for validation of data generated from clinical samples. Mouse studies may provide further space to delve into mechanistic details like cytokine regulatory network and cell types involved in IBD pathogenesis which is not possible with clinical samples. Therefore, we undertook studies in DSS induced colitis in mice to validate some of the clinical results.

### Disease phenotypes of DSS induced colitis in mouse and reversal with butyrate treatment

DSS colitis, despite shortcomings, has been of great use in understanding the pathophysiology of intestinal inflammation and, consequently, has informed the theoretical and clinical appreciation of active IBD (42). Here we induced colitis in mice with DSS and studied the pathogenies. The following phenotype was noted in colitis mouse and subsequent butyrate treatment: (a) significant decrease in colon length and body weight (Fig 8A, B, C) (b) 13.36-fold reduction of faecal butyrate as comparted to control mice [Fig 8D] and (c) significant decrease in IL-10 and increase in TNF-α and IFN-γ as compared to control [Fig8E] and (d) decrease in AUF1^p40^ mRNA and IL-27p28mRNA as compared to control [Fig8F, G]. Butyrate treatment to colitis mouse impelled significant reversal of above changes.

**Fig 8.**
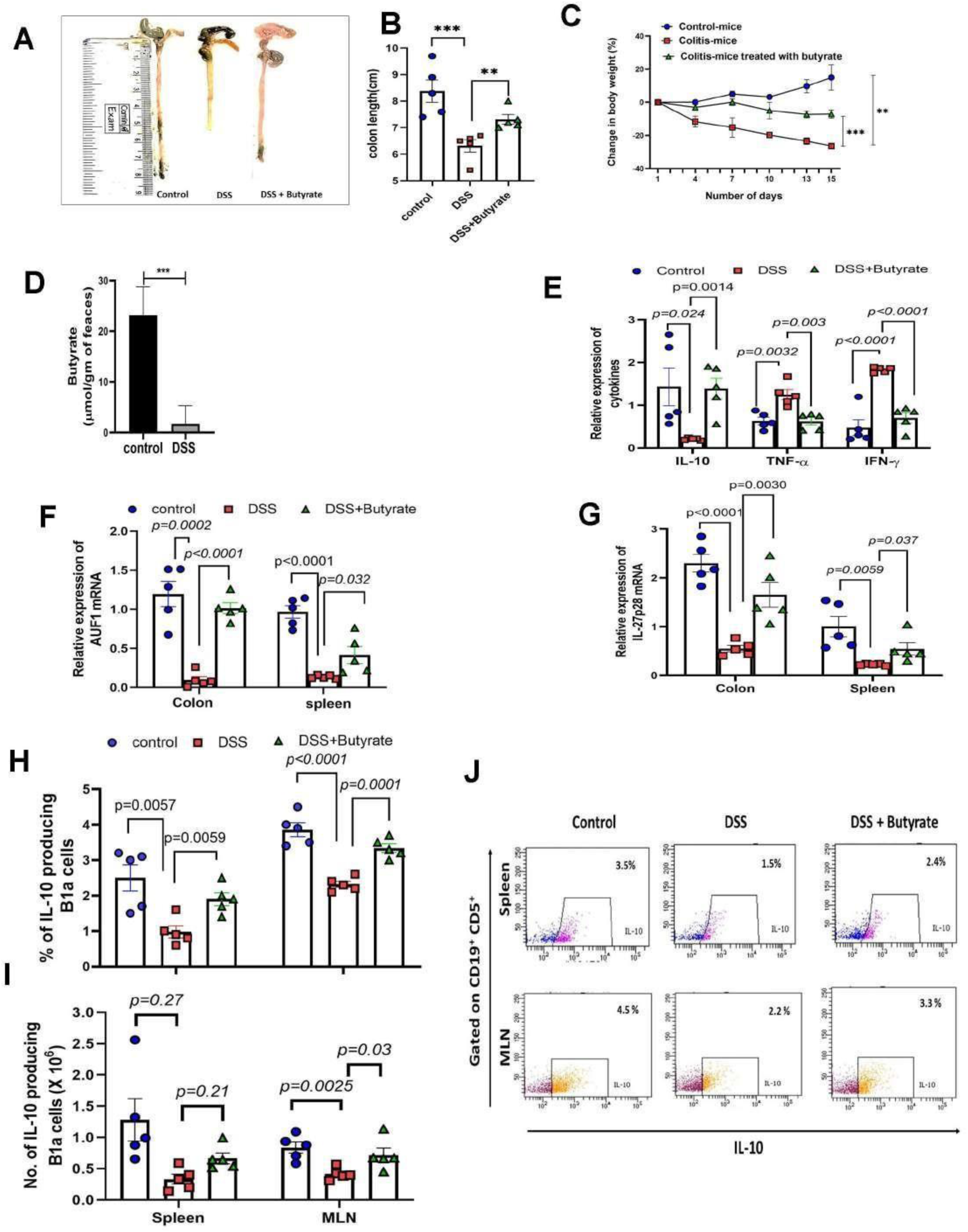
Butyrate protects DSS induced colitis in mice by increasing B10 polarization in spleen and mesenteric lymph nodes. Mice were randomly selected and divided into 3 groups (n=5 mice/group). First group was fed with 2.5% DSS for 14 days (colitis-mice), second group was fed with 2.5% DSS plus 5% butyrate with normal chow diet for 14 days (butyrate-colitis-mice). Third group received normal drinking water and food served as a control (control-mice). On day 15, mice were sacrificed and following parameters were analysed from all the mice from each group: colon length (A, B), % change in body weight (C), faecal butyrate measured by LC-MS and expressed in µmol/gm of faeces (D), expression of IL-10, TNF-a, and IFN-g in the colon tissue of control-mice, colitis-mice and butyrate-colitis-mice was measured by qPCR (E). Expression of AUF1 (F) and IL-27p28 (G) in the spleen and colon tissue of control-mice, colitis-mice and butyrate-colitis-mice was measured by qPCR. The cells from spleen and MLN isolated from each mouse were treated with monensin for 8 h. Thereafter the cells were stained with antibodies against CD19, CD5 and IL-10. The intracellular IL-10 positive cells were detected with their respective antibodies, gated on CD19^+^CD5^+^ cells. The percentage (H) and absolute number (I) of IL-10 producing CD19^+^CD5^+^ cells are represented. The representative dot plots showing IL-10 producing CD19^+^CD5^+^ cells (J). The gating and % CD19^+^CD5^+^ cells of spleen and MLN were shown in supplementary figure S5. Data represented of 3 independent experiments with n=5 mice/group. p value calculated by one-way ANOVA with post hoc Tukey’s test).

### Status of B1a cells in different secondary lymphoid organs in colitis mouse

Next, we went on to analyse percentage and number of IL-10 producing B1a cells not only in spleen but also in mesenteric lymph nodes (MLN) being in the proximity of the gut tissues. It was observed that frequency and absolute number of IL-10 producing B-1a cells were significantly decreased in colitis-mice regardless of origin and butyrate treatment increased their frequencies in low but significant measure [Fig 8H, I]. Notably the percentage of CD19^+^CD5+ cells (B1a cells) from spleen as well as in MLN remained unaltered irrespective of DSS and butyrate treatment (Fig S5).

### Adoptive transfer of butyrate treated B-1a from wild type mice but not from AUF1 knockdown mice conferred protection against DSS induced colitis

Since AUF-1 was decreased in clinical samples of colitis coupled with reduced IL-10 production and AUF-1 knock down mice display decrease in IL-10 producing B1a cells, we asked the question concerning the ability of butyrate treated B1a cells to attenuate DSS-induced intestinal injury and to establish role of AUF1 in roistering B1a to B10 polarization and buoying disease phenotype. The functional assay involved adoptive transfer of B1a cells either derived from S-MO treated mice or AUF1-MO (AUF1-KD) and adoptively transfer to colitis-mice. The experimental design is schematically represented in Figure 9A. B1a cells were sorted from either S-MO (wild-type) and AUF1- MO (AUF1-KD) mice were stimulated with 100 µM butyrate *in vitro* for 72 h. For convenience, B1a cells derived from S-MO or AUF1-MO defined as butyrate-B1a^wild^ and butyrate-B1a^KD^ cells respectively. The cells of butyrate-B1a^wild^ or butyrate-B1a^KD^ cells were injected into colitis-mice through tail vein. Adoptive transfer of butyrate-B1a^wild^ cells but not butyrate-B1a^KD^ cells to colitis-mice significantly reduced the severity of intestinal injury as observed by i) decrease in colon length (Fig 9 B, C), ii) significant decrease in TNF-α and IL-6 production in colon (Fig 9D) and iii) recovery from occult bleeding (Fig 9E).

**Fig 9:**
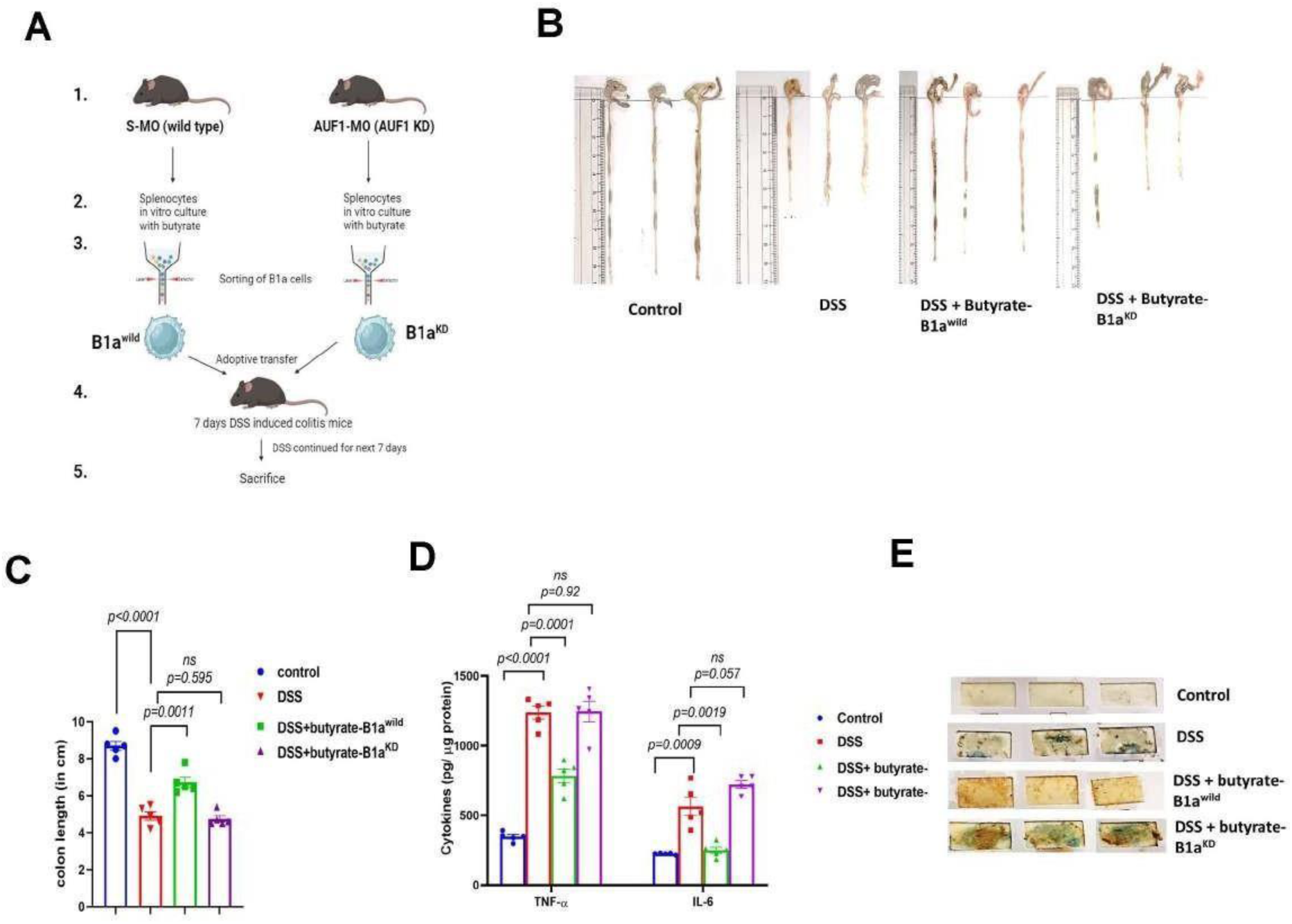
Butyrate treated B-1a from wild type mice but not the B-1a cells from AUF1 knockdown confer protection from DSS induced colitis. The experimental design is represented in five steps. A. Mice were injected with either S-MO or AUF1- MO as described in methodology. Splenocytes isolated from S-MO and AUF1-MO mice were then treated with butyrate (100 µM) for 72 h in vitro. Thereafter cells were stained with antibodies against CD19 and CD5 and gated and sorted by flowcytometry. The pre and post sort analysis is represented in Figure 6A and B. Butyrate treated CD19^+^CD5^+^ cells from S-MO mice and AUF1-MO mice were represented as butyrate-B1a^wild^ and butyrate-B1a-^KD^ cells respectively. Mice were fed with 2.5% DSS for 7 days and then injected with either 2X10^6^ cells butyrate-B1a^wild^ cells or 2 X 10^6^ cells butyrate- B1a^KD^ cells. The DSS administration was continued next 7 days. On day 15, the mice were sacrificed and following parameters were analysed: colon length (B) (C), expression of TNF-a and IL-6 in colon tissue as measured by ELISA (D) and occult bleeding (E). All experimented were repeated thrice with n=5 mice/group and represented as Mean± SE. p value calculated by one-way ANOVA with post hoc Tukey’s test).

## Discussion

Composing together series of experimental results, we dodged here linking butyrate with IL-10 producing B1a cells and identified the intermediate players and their mechanistic sequel that dangle in gut pathology of colitis. We chose to study with whole splenocytes or its gated population as the experiment demands. The spleen was of choice based on the report that IL-10 producing B cells are numerically higher in the spleen than in peritoneal cavity, lymph node (LN), Payer’s patch (PP) and gut associated lymphoid organs (37), (43). These numerically rare population of B cells producing IL-10 having unique functional program and suppresses inflammation are called “B10 cells” (37). Unlike acetate and propionate, butyrate has emerged as IL-10 producing short chain fatty acid from B1a cells but not in B2 cells, which has warranted a mechanistic look to understand how butyrate influences B10 polarization. The SCFA concentration used in the study was in tune with the report showing their physiological concentration in the peripheral blood in mice (44). The major regulatory pool of B cells come from B1a and B2 cells (45), therefore we have not considered B1b cells as it contributes mainly to adaptive responses (46).

Retroactively looking at the literature we found, IL-27, one of the IL-12 cytokine family members was shown to play an important role in IL-10 producing Treg cell development (33). Therefore, IL-27 induced B10 polarization was aptly looked into. We showed that B1a cells but not B2 cells produce IL- 10 upon IL-27 stimulation. We studied the expression of the two subunits of IL-27: IL-27p28, and EBI3 upon butyrate treatment and observed upregulation of IL-27p28 expression with butyrate compared to control. IL-27 is mostly produced from antigen presenting cells including macrophages, dendritic cells and also B1a cells but not from B2 cells (30). Therefore, in B1a cells IL-27 acts in autocrine as well as in paracrine manner. It is reported that B-1a cells express significantly higher IL-27 receptor (IL-27Rα) mRNA compared to B-2 cells (30). The lesser number of IL-27 receptor expression in B2 cell may explain its failure to induce IL-10 production upon stimulation with butyrate or IL-27. Furthermore, regardless of B cell lineage development, CD5-expressing B cells have a higher propensity for IL-10 production and this property has been associated with CD5 expression (47). To substantiate our finding that butyrate may increase IL-10 production in B1a cells through IL-27 upregulation, we blocked IL- 27 by using anti-IL-27 Ab in butyrate treated splenocytes. The significant reduction in the percentage of B10 cells from butyrate treated splenocytes upon IL-27 neutralization strengthens our hypothesis that butyrate increases the number of B10 cells by elevating IL-27 production. Earlier studies showed butyrate can increase IL-10 by increase in Blimp1 expression (48) and HDAC inhibition (49) whereas HDAC inhibitor inhibits Blimp-1-dependent gene repression (50). Since butyrate being an HDAC inhibitor (51), it may be speculated that it activates IL-10 by inhibiting Blimp-1 binding to its promoter. Interestingly, IL-27 is shown to activate IL-10 by c-Maf/ROR*γ*t/Blimp-1 signalling pathway (52). Therefore, earlier demonstration of butyrate mediated activation of IL-10 converges to the downstream of AUF1-IL-27 pathway.

IL-10 is regarded as a potent immune-suppressive cytokine with broad immunoregulatory functions (53). Thus, IL-10 suitably fulfils the profile of an effector molecule ‘downstream’ of IL-27 that exerts a general regulatory function of the B1a cells. Here, we showed that IL-27 not only increased expression of IL-10 but also promoted CD1d expression in B1a cells. Higher expression of CD1d is the hallmark of Breg phenotype (54). To further establish the anti-inflammatory function of butyrate stimulated B1a cells, we showed that the supernatant from B1a cells cultured in butyrate can reduce TNF-α production from LPS stimulated macrophage thereby endorsing the functional property of regulatory B cell in vitro. The cellular homeostasis of mRNA is not static and may vary in response to external stimuli through interacting with 3’-UTR. In addition to epigenetically regulating gene expression through its inhibition of HDAC activity, butyrate may also post-transcriptionally regulate the expression of various cytokine genes (27), (28), (29). To show the mechanism of regulation of IL-27 by butyrate, we undertook studies on the determining the decay rate of IL-27p28 mRNA in splenocytes. We found butyrate stabilizes IL- 27p28 mRNA increasing its half-life to 2.4-fold. TNF-α mRNA is reported to be destabilized by butyrate (27) therefore it was evident the half-life of TNF-α mRNA decreased upon butyrate treatment. Butyrate affects a number of other cellular functions which impact on gene expression, including RNA binding protein (RBP) mediated RNA stability (28), phosphorylation and ADP ribosylation of chromatin proteins and methylation of DNA (55). It is reported that butyrate regulates the expression of AU-rich element binding RBPs (56), (26) that can either accelerate or decelerate mRNA decay. The precise mechanism by which butyrate regulates these RBP expression is still not clear. AUF1which is reported to have dual role in mRNA stabilization as well as in decay by binding to the AU rich elements of mRNA (57) is expressed in spleen > thymus > intestine (58). AUF1 having 4 isoforms: p45, p42, p40 and p37 are important for B cell development (17), survival and expresses in B cells (59), macrophages (60) and epithelial cells (58). Consistent with the biological function of AUF1, our data demonstrate that AUF1 stabilizes IL-27 production by targeting p28 mRNA via direct binding to the 3′UTR of p28 mRNA. Although AUF1 reported to predominantly bind to the motifs located in the 3’UTR of the mRNA and decides the fate of mRNA (24), (61), it does not rule out the fact that it may also bind to other parts of p28 mRNA. Further studies are under way to address the different regions of AUF1 binding to p28 mRNA and also to identify the isoform(s) necessary for p28 mRNA stability. Oddly enough, another AU rich element binding RBP, Tristetraprolin (TTP) which is also upregulated by butyrate (62) is reported to destabilize IL-27p28 mRNA (63). TTP is abundant in liver and reproductive tissues but negligible in spleen where AUF1 is copious (64). Therefore, it can be speculated that AUF1 outpaces TTP on IL-27p28 mRNA regulation in the lymphoid tissues. AUF1 has pleiotropic targets, one of which is stabilizing IL-10 mRNA, by binding of p40 isoform of AUF1 (60). Refutably, we observed that IL-27 inhibition completely shuts down IL-10 production from B1a cells thereby contradicting the fact that AUF1 directly can increase IL-10 by stabilizing it’s mRNA. This may be explained by IL-27 mediated regulation of AUF1 in a positive feedback loop as evident from the western blot of AUF1 by inhibiting IL-27 expression using IL-27 morpholino (IL-27 MO) (Figure S6). To our surprise it was observed that blocking IL-27 by IL-27MO decreases the expression of AUF1 isoforms which is restored upon rIL-27 treatment. Further investigations are required to understand the phenomena of IL-27 mediated AUF1 regulation. Further investigations are required to understand the phenomena of IL-27 mediated AUF1 regulation.

To further establish the link between AUF-1 and IL-10 production in B1a cells, we knocked down AUF1 by deploying morpholino oligomer. Morpholino oligomers (MO) which are short single-stranded DNA analogues that are built upon a backbone of morpholine rings and are resistant to host enzymes present, a characteristic that makes them highly suitable for application in primary cells and *in vivo* (65), (26). Earlier we have used the AUF1 knock down mice model using MO to establish the role of AUF1 in cholesterol synthesis (26). Unlike a knockout, AUF1-MO mediated knock down causes partial knock down of AUF1 in organs like in kidney, heart (26), but nearly 100 % knock down in liver (26), colon and spleen. Therefore, warranting morpholino administration in mice a unique knockdown (KD) model to explore the role of AUF1 in B10 generation. The AUF1 KD mice showed low levels of IL-27 with decrease in the half-life of IL-27p28 mRNA compared to S-MO. Fundamentally, the process of mRNA decay is tightly linked to translation output (66) and we showed that absence of AUF1 represses the translation of IL27p28 mRNA. Reports showing the involvement of other trans factor(s) that are associated with translational repression with RBP mediated mRNA stability (67) provide fodder for further investigation of the trans factor (s) associated with AUF1 if any. The decrease in splenic B10 cells along with decrease in IL-27 in the spleen of AUF1 KD mice which butyrate failed to reverse, indicate that AUF1 is the master regulator of butyrate induced B10 polarization. Oddly enough, rIL-27 but not butyrate was perfectly capable of producing IL-10 from B1a cells in AUF-1-KD mice bringing credence to the notion that butyrate mediates its effect through IL-27. By aligning the observations, we can formulate the roster of molecular events as follows: butyrate-AUF1-IL-27p28-IL-10 in B1a cells. Since AUF1 regulates many cytokines gene expression the AUF1 knock out is reported to be susceptible to septicaemia and skeletal muscle wasting with impaired regeneration following injury (25), (68). Dialectic to the defects in post transcriptional regulation of the cytokines may be involved in human inflammatory diseases, we harnessed the pathways of IBD associated with AUF1 knockout cells by exploiting publicly-available microarray dataset by KEGG pathway analysis. Based on the alliance of IBD with AUF1 knockout we investigated the association of AUF1 and IL-27 in colon biopsy samples from IBD patients and non-IBD control. Significant decrease in AUF1 and the half-life of p28 mRNA in the colon biopsy of IBD-patients compared to control indicates an indispensable role of AUF1 in inflammation. Absence of AUF1 can also act as a colitogenic trigger as evident from the destructive colon architecture and increased inflammatory cytokines (TNF-α and IL-6) in AUF1 knock down mice compared to wild type control (Fig S7).

The reduction in AUF1 can be linked to IBD which is characterized by severe diminish in the IL-10 producing T and B regulatory cells (69, 70) and gut microbial dysbiosis resulting to decrease in butyrate (71), (72). As a surrogate of IBD patients we showed in DSS induced experimental colitis in mice reduction in butyrate and IL-10 producing B cells in MLN and spleen which was reversed upon butyrate treatment. Multiple studies have shown butyrate and rIL-27 administration ameliorates colitis in mice model (73), (74), (34). As there are very low frequency of B10 cells in gut associated lymphoid tissue (37) in wild type mice we preferred to analyse B10 cells from MLN considering its proximity to gut and also due to its sufficient availability in MLN (37). The decrease in B10 cells in colitis was aligned with decrease in AUF1 and IL-27 expression. Earlier it was shown that adoptive transfer with regulatory B1a cells decelerates colitis (75). To examine the importance of AUF1 in butyrate mediated B10 polarization in murine colitis, we isolated B1a cells from wild type mice and AUF1-KD mice to perform adoptive transfer in colitis-mice. We showed that butyrate treated B1a cells from wild type mice but not from AUF1KD mice exhibited therapeutic effects on colitis indicating B1a cells from wild type mice but not from AUF1 KD mice is polarized B10 upon butyrate treatment which attenuated inflammation. Therefore, we present quantifiable new data with the unique function of butyrate-driven B1a cells, offering an intriguing prospect for a colitis treatment.

In summary, this study unveils a novel regulatory pathway for IL-27 expression and B10 polarization mediated by AUF1, which contributes to the regulation of anti-inflammatory activity. It also indicates that butyrate induced AUF1 can dampen inflammation in DSS induced colitis through exploiting B10 polarization. Finally, we propose that butyrate driven B1a cells as a glimmer of new hope of therapeutic possibility against colitis.

## Materials and Methods

### Collection of colon biopsies and patient features

Eight active Ulcerative colitis (UC) patients as defined by Ungaro et al. (76) and seven patients who had colonoscopy and whose diagnosis for intestinal diseases were ruled out were enrolled for the study (Table 1). This study was approved by the ethics committee of National Institute of Cholera and Enteric Diseases (ICMR-NICED/IEC- BMHR/003/2023), and all methods were performed in accordance with the respective approval. Patients were informed in advance and gave written consent to sampling additional biopsies to ones that were taken for diagnostics during the routine endoscopic examination. For these, patients allowed usage for analysis of RNA and protein expression. From the Mayo endoscopic subscore is the part within the full Mayo score to access disease presentation during colonoscopy (77). Based on the aim of this study, patients with mild and moderate disease activity were unified as active (Table 1).

### Mice and animal ethics

C57BL/6 mice (4 weeks) were obtained from the animal facility of ICMR-NICED, Kolkata, India. All of the protocols for the study were approved by the Institutional Animal Ethics Committee of ICMR-NICED, Kolkata, India, (PRO/2001/-Nov 2023-26). Experiments have been carried out in accordance with the guidelines laid down by the committee for the purpose of control and supervision of experiments on animals (CPCSEA), Ministry of Environment and Forests, Government of India, New Delhi, India.

The daily consumption of food in mice was recorded every day by weighing the food given (food placed in the food receptacle at time zero) and food remaining (food left in the food receptacle 24 hours later). The cumulative food intake by each group (5 mice / group) per day was determined.

### Induction and Evaluation of DSS-Induced Intestinal Injury and calculation of Disease Activity Index (DAI)

2.5 % (w/v) of Dextran sodium sulphate (DSS) (molecular mass, 36 to 50 kDa) was dissolved in purified water and administered to mice for 14 days (78). In some experiments, mice were fed with 5% butyrate (w/w) mixed with food pellet from the day of experiment initiation till it ended (26). The DAI is the combined score of weight loss compared to initial weight, stool consistency, and bleeding. Scores are defined as follows: weight loss: 0 (no loss), 1 (1-5%), 2 (5-10%), 3 (10-20%), and 4 (>20%); stool consistency: 0 (normal), 2 (loose stool), and 4 (diarrhoea); and bleeding: 0 (no blood), 1 (Hemoccult positive), 2 (Hemoccult positive and visual pellet bleeding), and 4 (gross bleeding, blood around anus) (78).

### Histological Analysis

The mice were sacrificed 5 days after the induction of intestinal injury. Colon samples were removed and segments were fixed in 10% buffered formalin. After paraffin embedding, 5-μm-thick sections were cut and stained with H&E. Histological scoring was based on a previously described method (79). Briefly, H&E-stained cross sections of the descending colon tissue were scored microscopically in a blinded fashion on a scale from 0 to 4, based on the following histological criteria: 0, no change from normal tissue; 1, low level of inflammation with scattered infiltrating mononuclear cells (foci, 1 to 2); 2, moderate inflammation with multiple foci; 3, high level of inflammation with increased vascular density and marked wall thickening; and 4, maximal severity of inflammation with transmural leukocyte infiltration and loss of goblet cells. An average of four fields of view per colon was evaluated for each mouse. These scores were averaged for each group and recorded as the histopathological score.

### Intracellular staining

Single-cell suspension of splenocytes from mice was prepared as described previously (80). The cells were treated as indicated in the experiments and then cultured in the presence of 10 µg/ml Monensin for 8 h. Cells were stained by monoclonal fluorescent tagged antibodies either against CD3, CD4, or CD19, CD5 or CD 23 or CD1d fixed and permeated using Cytofix/Cytoperm, followed by intracellular staining with anti-IL-10 antibody. Details of the antibodies are given in the supplement.

### Cell Sorting and Adoptive Transfers

Spleen cells from scramble-MO and AUF1-MO mice were isolated and treated with 100 µM butyrate for 72 h at 37°C in humified CO_2_ incubator. After incubation, B1a cells (CD19^+^CD5^+^ B cells) from butyrate treated spleen cells of scramble-MO and AUF1-MO mice were purified using a flow cytometer (FACSAria Fusion; Becton Dickinson), with purities of approximately 85% to 95%. The 2 × 10^6^ cells in 200 µl was transferred by intravenous injection into 7 days post DSS fed mice. On day 7^th^ of post injection, the mice were sacrificed and colon were collected, length of the colon was measured and further processed for histology and cytokine analysis.

### GMO-PMO treatment to knock down AUF1

The GMO-PMO synthesis and its administration in mice was performed as described earlier (26). Briefly, The GMO-PMO was dissolved in saline to make a concentration of 1 mM which was equivalent to 8.2 mg/ml. In vivo knock down of AUF1 was carried out in mice by injecting AUF1 GMO-PMO (AUF1-MO) through tail vein on day 1 and day 7 at a dose of 3 mg/kg body weight. Scramble GMO-PMO treated animals are called as S-MO controls. Animals were sacrificed after day 14 and tissue were collected for further analysis.

Sequence of AUF1-MO = 5’-TCCGAACTGCTCCTCCGACATAGTG-3’

Sequence of IL-27-MO = 5’ CAAGGTCTCCTGTCACCTGGCCCAT3’

Sequence of S-MO **=** 5’-TCGCAACTCGTCCTCCAGCATAGTG-3’

### Cloning of IL-27p28 3’UTR

Mouse IL-27 p28 3′UTR plasmid was cloned by inserting p28 3′UTR into MCS of the pEGFP-C1 vector (Clontech, California US) between XhoI and KpnI sites. Primers used for amplification of p28 3′UTR from genomic DNA of splenocytes were CCGCTCGAGTTCTAGACACCTAGCTTCAAGCCCTATGG (sense); and CGGGGTACCGGCCGGCCCGGGCTGGATGGCTTTATTA (anti-sense). All plasmid DNA were prepared with QIAGEN Endo-free Mini-Prep kits.

### Transfection

RAW264.7 cells were plated in 6-well culture plates at the density of 1×10^6^ cells/well and cultured in 2 ml serum-free medium for 24 h to 80% confluence. Transfection was performed with Lipofectamine 2000 according to the protocol recommended by the manufacturer. Briefly, 25 μl Opti-MEM medium was used to dilute 1.0 μl lipofectamine and 0.5 μg plasmid or 0.27 μg siRNA, and equal volume of the plasmid or siRNA and lipofectamine was mixed at room temperature for 15 min. Cells were transfected by adding 100 μl of the Opti-MEM medium containing lipofectamine and plasmid or siRNA at 37° C for 6 h, and then cells were grown at DMEM containing 10% FCS.

## Fluorescence microscopy

RAW264.7 cells grown on glass cover slips were transfected with either pEGFP-IL-27p28-3′UTR or p- EGFP for 24 h. After washing with PBS, the cells were treated with 1 µg/ml Hoechst 33342 for 5 min at room temperature and washed again with PBS three times. Fluorescence images were captured with Carl Zeiss microscope equipped with a CCD camera controlled with ZEN software (Carl Zeiss, Gottingen, Germany). The fluorescence intensity was quantified and expressed as corrected total cell fluorescence (CTCF). CTCF = integrated density – (area of selected cells X mean fluorescence of background readings) (81).

### Determination of mRNA half-life

Splenocytes were challenged with Actinomycin D (5µg/ml) for increasing periods of time. Total RNA was then extracted and subjected to qPCR. The one-phase exponential decay curve analysis (GraphPad Prism 9) was used to assess the mRNA decay kinetics, considering the values at time 0 as 1. The time corresponding to 50% remaining mRNA was considered as mRNA half-life (t_1/2_) (63).

### RNA immunoprecipitation

Proteins extracted from whole splenocytes were incubated with beads pre-coated with anti-AUF1 antibody and control IgG. After washing three times, RNA was extracted from Beads, and reverse- transcripted into cDNA, followed by detecting p28 and TNF α mRNA by qPCR (63).

### Quantitative reverse transcription-PCR

Total RNA was extracted from B cells or colon tissue using Trizol reagent and RNA quantification was performed on Nanodrop 2000. RNA (0.5–1μg) was reverse transcribed using the cDNA Synthesis Kit according to the manufacturer’s instructions. The qPCR reaction was performed using the SYBR Green Kit and on QuantStudio 5/7 (Applied Biosystems, Massachusetts, USA). Primers were designed and synthesized with the sequences listed below. All primers were purchased from IDT (Lowa, USA) (Supplementary Table).

### Polysome analysis

Ribosomal fractions were obtained as described earlier (75). In brief, spleen cells were homogenized in polysome lysis buffer containing cycloheximide (0.1 mg/ml) and cytosolic extract was obtained by centrifugation at 10000g for 20 min. The extract was overlayed on a 10–50% (w/v) sucrose gradient and centrifuged at 100000g for 4 h in ultracentrifuge (Thermo, Massechusetts, USA). Fractions were collected manually and absorbance at 254 nm was measured. RNA was isolated from the fractions by phenol-chloroform extraction and ethanol precipitation.

### Western blot

Tissue protein or cell lysate were extracted in RIPA Lysis buffer. Protein concentration was measured using Pierce BCA Protein Assay Kit. Proteins (50 μg/lane) were separated by using SDS-PAGE on 10% gel under reducing condition and electro transferred to PVDF membrane in a transferred buffer (25mM Tris-HCl, 150mM Glycine, 20% Methanol). Membranes were blocked at room temperature with 5% non fat skim milk in TBS for 2 hours, and then incubated with primary antibody against specific protein. The membranes were incubated either with the horseradish peroxidase (HRP)-conjugated secondary antibodies or Alkaline phosphatase (AP)-conjugated antibodies at 37° C for 1 h. For HRP-conjugated antibody treatment, SuperSignal West Pico chemiluminescent substrate kit (Thermo) was used to visualize the blotting results. The blots were imaged with Fluor Chem R system (ProteinSimple, San Jose, CA, USA) (76). For AP-conjugated antibody treatment, BCIP-NBT substrate kit (Thermo) was used to visualize the blotting results (77).

### KEGG pathway analysis

The data (GSE138621) FastQ files were obtained from NCBI-SRA, specifically processing samples belonging to the specified groups (AUF1-knockdown WS1 cell line vs WS1 wild type cell control). Initially, the samples underwent quality filtering using trim_galore. Next, quality-controlled samples were aligned to the human reference genome using the STAR aligner (78). Subsequently, featureCounts (79) was employed on the generated BAM files to produce a count table. This count table underwent further analysis using the DESeq2 (80) package in R. Additionally, KEGG pathway over-representation analysis was conducted separately for upregulated and downregulated genes, based on specific criteria (log2fold change <= -0.5 or >= 0.5 and a p-value less than 0.05), utilizing the clusterProfiler (81) package.

### Measurement of butyrate in faeces by LC-MS

Faecal content of each mouse (10-50 mg wet weight) was dissolved in distilled water at 1:10 (w/v) and homogenized in a Dounce homogenizer. Then, the content was centrifuged at 10,000g for 10 m and the supernatant was filtered through 0.45um syringe filter. The supernatant was further diluted in LC-MS grade distilled water up to a final volume of 1 ml and were subjected to LC-MS analysis for SCFA as described by Cheng et al (82). In brief, a calibration curve was generated by using varying calibration of internal standards of butyrate from 2 µmoles to 50 µmoles. The LCMS separation was performed on the Agilent 1290 Infinity LC system which was coupled to Agilent 6545 Accurate-Mass Quadrupole Time-of Flight (QTOF) with Agilent Jet Stream Thermal Gradient Technology with electrospray ionization (ESI) source. The suitable MS parameters were optimised and the high-resolution mass spectra were obtained by performing the analysis in negative ionisation mode. The Chromatographic separation was achieved on Agilent ZORBAX SB- C18 column (2.1 × 100 mm, 1.8 µm) as stationary phase. The mobile phase consisted of a linear gradient of 0.1% (v/v) aqueous formic acid (A) and Methanol (B): 0–1.0 min, 0 % B (v/v); 1.0–3.0 min, 0 30% B (v/v); 3.0–6.0 min, 30-100% B (v/v); 6.0–12.0 min, 100% B (v/v); 12.50–15.0 min, 0% B. The column was reconditioned for 5 min before next injection. 0.2 mL/min flow rate and a varying injection volume was used for analysis. The UPLC system assembled with a Diode array detector (DAD) and an auto sampler. The peak area of each SCFA was used to calculate the amount of SCFA present which was further normalized by the injection volume. The data was further represented as the amount of SCFA present per gram of faeces.

### Statistics

All graphs and statistical significance were calculated using GraphPad Prism 9.00 (GraphPad, San Diego). For mouse studies, we chose a sample size of five mice per group. Unpaired two-tailed Student’s t test was performed for comparison between two groups. One-way ANOVA (Nonparametric) with Tukey (Compare all pairs of columns) or Kruskal–Walls test with Dunn’s Multiple Comparison was used for comparison among multiple groups. Data shown are mean+SE. of at least three independent experiments. *p < 0.05; **p < 0.01; ***p < 0.001 between indicated groups. Curve fitting, the mathematical modelling, was done with Origin Lab data analysis.

## Acknowledgements

We acknowledge the support of Director, ICMR-NICED, Kolkata for carrying out the study. Our deepest gratitude to Prof Syamal Roy (CSIR-IICB, Kolkata), Dr. Dipsikha Chakravarty (IISc, Bangalore), Dr. Kaushik Roy (University of Utah, USA) for helpful discussions and suggestions for the manuscript. We thank Dr. Tapas Biswas and Dr. Debalina Sinha for initiating the idea that IL-27 may induce IL-10 production from B1a cells. We thank Dr. Rajib Sarkar (SSKM Hospital, Kolkata) for clinical samples, Dr Rahul Gajbhiye (NIPER, Mohali) for butyrate measurement by LC-MS, Dr. Anupam Gautam (University of Tubingen, Germany) for KEGG pathway analysis and Payel (CSIR-IICB, Kolkata) and Bhaswati Tarafdar (Tata Cancer Centre, Kolkata) for FACS analysis. We acknowledge Narayan Mondal, Aranya Karmakar, Manik Barik, Anita Goswami and Usha Hansdah for animal maintenance and sample collection. AM, SS, OD, DR are recipients of fellowship from the CSIR, DST and UGC respectively.

## Author contributions

AM performed majority of the experiments and analysed data. OD showed that AUF1 knockdown mice develops colitis and performed experiments on clinical samples. DR and SAC performed the cloning. AD, SS, SA performed experiments. RS and MY provided clinical samples and helped in diagnosis of the disease by colonoscopy. SS synthesized the morpholino. MB conceptualized, designed, supervised, arranged funding and wrote the manuscript.

## Data availability statement

All data generated in this study are included in this article and supplementary information or are available from the corresponding author on reasonable request.

## References

1. Golpour F, Abbasi-Alaei M, Babaei F, Mirzababaei M, Parvardeh S, Mohammadi G, et al. Short chain fatty acids, a possible treatment option for autoimmune diseases. Biomed Pharmacother. 2023;163:114763.

2. Luhrs H, Gerke T, Muller JG, Melcher R, Schauber J, Boxberge F, et al. Butyrate inhibits NF-kappaB activation in lamina propria macrophages of patients with ulcerative colitis. Scand J Gastroenterol. 2002;37(4):458–66.

3. Scheppach W, Sommer H, Kirchner T, Paganelli GM, Bartram P, Christl S, et al. Effect of butyrate enemas on the colonic mucosa in distal ulcerative colitis. Gastroenterology. 1992;103(1):51–6.

4. Goetz M, Atreya R, Ghalibafian M, Galle PR, Neurath MF. Exacerbation of ulcerative colitis after rituximab salvage therapy. Inflamm Bowel Dis. 2007;13(11):1365–8.

5. Olson TS, Bamias G, Naganuma M, Rivera-Nieves J, Burcin TL, Ross W, et al. Expanded B cell population blocks regulatory T cells and exacerbates ileitis in a murine model of Crohn disease. J Clin Invest. 2004;114(3):389–98.

6. Sun X, Huang Y, Zhang YL, Qiao D, Dai YC. Research advances of vasoactive intestinal peptide in the pathogenesis of ulcerative colitis by regulating interleukin-10 expression in regulatory B cells. World J Gastroenterol. 2020;26(48):7593–602.

7. Fillatreau S, Sweenie CH, McGeachy MJ, Gray D, Anderton SM. B cells regulate autoimmunity by provision of IL-10. Nat Immunol. 2002;3(10):944–50.

8. Tsuzuki Y, Shiomi R, Ashitani K, Miyaguchi K, Osaki A, Ohgo H, et al. Rituximab- induced Ileocolitis in a Patient with Gastric MALToma: A Case Report and Literature Review. Intern Med. 2021;60(5):731–8.

9. Chan CC. Couching for cataract in China. Surv Ophthalmol. 2010;55(4):393–8.

10. Obesity 2009. Abstracts of the 27th Annual Scientific Meeting of The Obesity Society, October 24-28, 2009, Washington, DC, USA. Obesity (Silver Spring). 2009;17 Suppl 2:S1–373.

11. Baumgarth N. The double life of a B-1 cell: self-reactivity selects for protective effector functions. Nat Rev Immunol. 2011;11(1):34–46.

12. Scheiner R, Toteva A, Reim T, Sovik E, Barron AB. Differences in the phototaxis of pollen and nectar foraging honey bees are related to their octopamine brain titers. Front Physiol. 2014;5:116.

13. Berland R, Wortis HH. Origins and functions of B-1 cells with notes on the role of CD5. Annu Rev Immunol. 2002;20:253–300.

14. Baumgarth N. A Hard(y) Look at B-1 Cell Development and Function. J Immunol. 2017;199(10):3387–94.

15. Mahajan VS, Mattoo H, Sun N, Viswanadham V, Yuen GJ, Allard-Chamard H, et al. B1a and B2 cells are characterized by distinct CpG modification states at DNMT3A- maintained enhancers. Nat Commun. 2021;12(1):2208.

16. White EJ, Matsangos AE, Wilson GM. AUF1 regulation of coding and noncoding RNA. Wiley Interdiscip Rev RNA. 2017;8(2).

17. Sadri N, Lu JY, Badura ML, Schneider RJ. AUF1 is involved in splenic follicular B cell maintenance. BMC Immunol. 2010;11:1.

18. AlAhmari MM, Al-Khalaf HH, Al-Mohanna FH, Ghebeh H, Aboussekhra A. AUF1 promotes stemness in human mammary epithelial cells through stabilization of the EMT transcription factors TWIST1 and SNAIL1. Oncogenesis. 2020;9(8):70.

19. Zou T, Rao JN, Liu L, Xiao L, Yu TX, Jiang P, et al. Polyamines regulate the stability of JunD mRNA by modulating the competitive binding of its 3’ untranslated region to HuR and AUF1. Mol Cell Biol. 2010;30(21):5021–32.

20. Lafon I, Carballes F, Brewer G, Poiret M, Morello D. Developmental expression of AUF1 and HuR, two c-myc mRNA binding proteins. Oncogene. 1998;16(26):3413–21.

21. Skriner K, Hueber W, Suleymanoglu E, Hofler E, Krenn V, Smolen J, et al. AUF1, the regulator of tumor necrosis factor alpha messenger RNA decay, is targeted by autoantibodies of patients with systemic rheumatic diseases. Arthritis Rheum. 2008;58(2):511–20.

22. Raineri I, Wegmueller D, Gross B, Certa U, Moroni C. Roles of AUF1 isoforms, HuR and BRF1 in ARE-dependent mRNA turnover studied by RNA interference. Nucleic Acids Res. 2004;32(4):1279–88.

23. Khabar KS. Post-transcriptional control during chronic inflammation and cancer: a focus on AU-rich elements. Cell Mol Life Sci. 2010;67(17):2937–55.

24. Gratacos FM, Brewer G. The role of AUF1 in regulated mRNA decay. Wiley Interdiscip Rev RNA. 2010;1(3):457–73.

25. Lu JY, Sadri N, Schneider RJ. Endotoxic shock in AUF1 knockout mice mediated by failure to degrade proinflammatory cytokine mRNAs. Genes Dev. 2006;20(22):3174–84.

26. Das O, Kundu J, Ghosh A, Gautam A, Ghosh S, Chakraborty M, et al. AUF-1 knockdown in mice undermines gut microbial butyrate-driven hypocholesterolemia through AUF-1-Dicer-1-mir-122 hierarchy. Front Cell Infect Microbiol. 2022;12:1011386.

27. Fukae J, Amasaki Y, Yamashita Y, Bohgaki T, Yasuda S, Jodo S, et al. Butyrate suppresses tumor necrosis factor alpha production by regulating specific messenger RNA degradation mediated through a cis-acting AU-rich element. Arthritis Rheum. 2005;52(9):2697–707.

28. Torun A, Enayat S, Sheraj I, Tuncer S, Ulgen DH, Banerjee S. Butyrate mediated regulation of RNA binding proteins in the post-transcriptional regulation of inflammatory gene expression. Cell Signal. 2019;64:109410.

29. Zheng L, Kelly CJ, Battista KD, Schaefer R, Lanis JM, Alexeev EE, et al. Microbial-Derived Butyrate Promotes Epithelial Barrier Function through IL-10 Receptor-Dependent Repression of Claudin-2. J Immunol. 2017;199(8):2976–84.

30. Choi JK, Yu CR, Bing SJ, Jittayasothorn Y, Mattapallil MJ, Kang M, et al. IL-27- producing B-1a cells suppress neuroinflammation and CNS autoimmune diseases. Proc Natl Acad Sci U S A. 2021;118(47).

31. Lin CH, Wu CJ, Cho S, Patkar R, Lin LL, Chen MC, et al. Selective IL-27 production by intestinal regulatory T cells permits gut-specific regulation of Th17 immunity. bioRxiv. 2023.

32. Kang M, Yadav MK, Mbanefo EC, Yu CR, Egwuagu CE. IL-27-containing exosomes secreted by innate B-1a cells suppress and ameliorate uveitis. Front Immunol. 2023;14:1071162.

33. Murugaiyan G, Mittal A, Lopez-Diego R, Maier LM, Anderson DE, Weiner HL. IL-27 is a key regulator of IL-10 and IL-17 production by human CD4+ T cells. J Immunol. 2009;183(4):2435–43.

34. McLean MH, Andrews C, Hanson ML, Baseler WA, Anver MR, Senkevitch E, et al. Interleukin-27 Is a Potential Rescue Therapy for Acute Severe Colitis Through Interleukin-10- Dependent, T-Cell-Independent Attenuation of Colonic Mucosal Innate Immune Responses. Inflamm Bowel Dis. 2017;23(11):1983–95.

35. Branchett WJ, Lloyd CM. Regulatory cytokine function in the respiratory tract. Mucosal Immunol. 2019;12(3):589–600.

36. Min B, Kim D, Feige MJ. IL-30(dagger) (IL-27A): a familiar stranger in immunity, inflammation, and cancer. Exp Mol Med. 2021;53(5):823–34.

37. Lykken JM, Candando KM, Tedder TF. Regulatory B10 cell development and function. Int Immunol. 2015;27(10):471–7.

38. Liu H, Rohowsky-Kochan C. Interleukin-27-mediated suppression of human Th17 cells is associated with activation of STAT1 and suppressor of cytokine signaling protein 1. J Interferon Cytokine Res. 2011;31(5):459–69.

39. Kumar P, Rajasekaran K, Nanbakhsh A, Gorski J, Thakar MS, Malarkannan S. IL-27 promotes NK cell effector functions via Maf-Nrf2 pathway during influenza infection. Sci Rep. 2019;9(1):4984.

40. Oleinika K, Rosser EC, Matei DE, Nistala K, Bosma A, Drozdov I, et al. CD1d- dependent immune suppression mediated by regulatory B cells through modulations of iNKT cells. Nat Commun. 2018;9(1):684.

41. Al-Khalaf HH, Aboussekhra A. AUF1 positively controls angiogenesis through mRNA stabilization-dependent up-regulation of HIF-1alpha and VEGF-A in human osteosarcoma. Oncotarget. 2019;10(47):4868–79.

42. Eichele DD, Kharbanda KK. Dextran sodium sulfate colitis murine model: An indispensable tool for advancing our understanding of inflammatory bowel diseases pathogenesis. World J Gastroenterol. 2017;23(33):6016–29.

43. Burger C, Vitetta ES. The response of B cells in spleen, Peyer’s patches, and lymph nodes to LPS and IL-4. Cell Immunol. 1991;138(1):35–43.

44. Fuller M, Priyadarshini M, Gibbons SM, Angueira AR, Brodsky M, Hayes MG, et al. The short-chain fatty acid receptor, FFA2, contributes to gestational glucose homeostasis. Am J Physiol Endocrinol Metab. 2015;309(10):E840–51.

45. Rosser EC, Mauri C. Regulatory B cells: origin, phenotype, and function. Immunity. 2015;42(4):607–12.

46. Haas KM, Poe JC, Steeber DA, Tedder TF. B-1a and B-1b cells exhibit distinct developmental requirements and have unique functional roles in innate and adaptive immunity to S. pneumoniae. Immunity. 2005;23(1):7–18.

47. O’Garra A, Chang R, Go N, Hastings R, Haughton G, Howard M. Ly-1 B (B-1) cells are the main source of B cell-derived interleukin 10. Eur J Immunol. 1992;22(3):711–7.

48. Sun M, Wu W, Chen L, Yang W, Huang X, Ma C, et al. Microbiota-derived short-chain fatty acids promote Th1 cell IL-10 production to maintain intestinal homeostasis. Nat Commun. 2018;9(1):3555.

49. Foh B, Buhre JS, Lunding HB, Moreno-Fernandez ME, Konig P, Sina C, et al. Microbial metabolite butyrate promotes induction of IL-10+IgM+ plasma cells. PLoS One. 2022;17(3):e0266071.

50. Yu J, Angelin-Duclos C, Greenwood J, Liao J, Calame K. Transcriptional repression by blimp-1 (PRDI-BF1) involves recruitment of histone deacetylase. Mol Cell Biol. 2000;20(7):2592–603.

51. Steliou K, Boosalis MS, Perrine SP, Sangerman J, Faller DV. Butyrate histone deacetylase inhibitors. Biores Open Access. 2012;1(4):192–8.

52. Newman JH. Lung vascular injury. Chest. 1988;93(3 Suppl):139S–46S.

53. Iyer SS, Cheng G. Role of interleukin 10 transcriptional regulation in inflammation and autoimmune disease. Crit Rev Immunol. 2012;32(1):23–63.

54. Karim MR, Wang YF. Phenotypic identification of CD19(+)CD5(+)CD1d(+) regulatory B cells that produce interleukin 10 and transforming growth factor beta(1) in human peripheral blood. Arch Med Sci. 2019;15(5):1176–83.

55. Mathew OP, Ranganna K, Mathew J, Zhu M, Yousefipour Z, Selvam C, et al. Cellular Effects of Butyrate on Vascular Smooth Muscle Cells are Mediated through Disparate Actions on Dual Targets, Histone Deacetylase (HDAC) Activity and PI3K/Akt Signaling Network. Int J Mol Sci. 2019;20(12).

56. Maclean KN, McKay IA, Bustin SA. Differential effects of sodium butyrate on the transcription of the human TIS11 family of early-response genes in colorectal cancer cells. Br J Biomed Sci. 1998;55(3):184–91.

57. White EJ, Brewer G, Wilson GM. Post-transcriptional control of gene expression by AUF1: mechanisms, physiological targets, and regulation. Biochim Biophys Acta. 2013;1829(6-7):680–8.

58. Caviglia JM, Li LO, Wang S, DiRusso CC, Coleman RA, Lewin TM. Rat long chain acyl-CoA synthetase 5, but not 1, 2, 3, or 4, complements Escherichia coli fadD. J Biol Chem. 2004;279(12):11163–9.

59. Diaz-Munoz MD, Bell SE, Turner M. Deletion of AU-rich elements within the Bcl2 3’UTR reduces protein expression and B cell survival in vivo. PLoS One. 2015;10(2):e0116899.

60. Sarkar S, Sinsimer KS, Foster RL, Brewer G, Pestka S. AUF1 isoform-specific regulation of anti-inflammatory IL10 expression in monocytes. J Interferon Cytokine Res. 2008;28(11):679–91.

61. Zucconi BE, Wilson GM. Assembly of functional ribonucleoprotein complexes by AU-rich element RNA-binding protein 1 (AUF1) requires base-dependent and -independent RNA contacts. J Biol Chem. 2013;288(39):28034–48.

62. Zheng XT, Xiao XQ, Dai JJ. Corrigendum to “Sodium butyrate down-regulates tristetraprolin-mediated cyclin B1expression independent of the formation of processing bodies” [Int. J. Biochem. Cell Biol. 69 (2015) 241–248]. Int J Biochem Cell Biol. 2016;74:161.

63. Wang Q, Ning H, Peng H, Wei L, Hou R, Hoft DF, et al. Tristetraprolin inhibits macrophage IL-27-induced activation of antitumour cytotoxic T cell responses. Nat Commun. 2017;8(1):867.

64. Lu JY, Schneider RJ. Tissue distribution of AU-rich mRNA-binding proteins involved in regulation of mRNA decay. J Biol Chem. 2004;279(13):12974–9.

65. Das U, Kundu J, Shaw P, Bose C, Ghosh A, Gupta S, et al. Self-transfecting GMO-PMO chimera targeting Nanog enable gene silencing in vitro and suppresses tumor growth in 4T1 allografts in mouse. Mol Ther Nucleic Acids. 2023;32:203–28.

66. Roy B, Jacobson A. The intimate relationships of mRNA decay and translation. Trends Genet. 2013;29(12):691–9.

67. Kumar R, Poria DK, Ray PS. RNA-binding proteins La and HuR cooperatively modulate translation repression of PDCD4 mRNA. J Biol Chem. 2021;296:100154.

68. Chenette DM, Cadwallader AB, Antwine TL, Larkin LC, Wang J, Olwin BB, et al. Targeted mRNA Decay by RNA Binding Protein AUF1 Regulates Adult Muscle Stem Cell Fate, Promoting Skeletal Muscle Integrity. Cell Rep. 2016;16(5):1379–90.

69. Oka A, Ishihara S, Mishima Y, Tada Y, Kusunoki R, Fukuba N, et al. Role of regulatory B cells in chronic intestinal inflammation: association with pathogenesis of Crohn’s disease. Inflamm Bowel Dis. 2014;20(2):315–28.

70. Yamada A, Arakaki R, Saito M, Tsunematsu T, Kudo Y, Ishimaru N. Role of regulatory T cell in the pathogenesis of inflammatory bowel disease. World J Gastroenterol. 2016;22(7):2195–205.

71. Parada Venegas D, De la Fuente MK, Landskron G, Gonzalez MJ, Quera R, Dijkstra G, et al. Short Chain Fatty Acids (SCFAs)-Mediated Gut Epithelial and Immune Regulation and Its Relevance for Inflammatory Bowel Diseases. Front Immunol. 2019;10:277.

72. Haneishi Y, Furuya Y, Hasegawa M, Picarelli A, Rossi M, Miyamoto J. Inflammatory Bowel Diseases and Gut Microbiota. Int J Mol Sci. 2023;24(4).

73. Dou X, Gao N, Yan D, Shan A. Sodium Butyrate Alleviates Mouse Colitis by Regulating Gut Microbiota Dysbiosis. Animals (Basel). 2020;10(7).

74. Li G, Lin J, Zhang C, Gao H, Lu H, Gao X, et al. Microbiota metabolite butyrate constrains neutrophil functions and ameliorates mucosal inflammation in inflammatory bowel disease. Gut Microbes. 2021;13(1):1968257.

75. Yanaba K, Yoshizaki A, Asano Y, Kadono T, Tedder TF, Sato S. IL-10-producing regulatory B10 cells inhibit intestinal injury in a mouse model. Am J Pathol. 2011;178(2):735–43.

76. Ungaro R, Mehandru S, Allen PB, Peyrin-Biroulet L, Colombel JF. Ulcerative colitis. Lancet. 2017;389(10080):1756–70.

77. Schroeder KW, Tremaine WJ, Ilstrup DM. Coated oral 5-aminosalicylic acid therapy for mildly to moderately active ulcerative colitis. A randomized study. N Engl J Med. 1987;317(26):1625–9.

78. Kim JJ, Shajib MS, Manocha MM, Khan WI. Investigating intestinal inflammation in DSS-induced model of IBD. J Vis Exp. 2012(60).

79. Erben U, Loddenkemper C, Doerfel K, Spieckermann S, Haller D, Heimesaat MM, et al. A guide to histomorphological evaluation of intestinal inflammation in mouse models. Int J Clin Exp Pathol. 2014;7(8):4557–76.

80. Guha R, Gupta D, Rastogi R, Vikram R, Krishnamurthy G, Bimal S, et al. Vaccination with leishmania hemoglobin receptor-encoding DNA protects against visceral leishmaniasis. Sci Transl Med. 2013;5(202):202ra121.

81. Bora P, Gahurova L, Masek T, Hauserova A, Potesil D, Jansova D, et al. p38-MAPK-mediated translation regulation during early blastocyst development is required for primitive endoderm differentiation in mice. Commun Biol. 2021;4(1):788.

82. Panda AC, Martindale JL, Gorospe M. Polysome Fractionation to Analyze mRNA Distribution Profiles. Bio Protoc. 2017;7(3).

83. Abdelmohsen K, Tominaga-Yamanaka K, Srikantan S, Yoon JH, Kang MJ, Gorospe M. RNA-binding protein AUF1 represses Dicer expression. Nucleic Acids Res. 2012;40(22):11531–44.

84. Blake MS, Johnston KH, Russell-Jones GJ, Gotschlich EC. A rapid, sensitive method for detection of alkaline phosphatase-conjugated anti-antibody on Western blots. Anal Biochem. 1984;136(1):175–9.

85. Dobin A, Davis CA, Schlesinger F, Drenkow J, Zaleski C, Jha S, et al. STAR: ultrafast universal RNA-seq aligner. Bioinformatics. 2013;29(1):15–21.

86. Liao Y, Smyth GK, Shi W. featureCounts: an efficient general purpose program for assigning sequence reads to genomic features. Bioinformatics. 2014;30(7):923–30.

87. Love MI, Huber W, Anders S. Moderated estimation of fold change and dispersion for RNA-seq data with DESeq2. Genome Biol. 2014;15(12):550.

88. Wu T, Hu E, Xu S, Chen M, Guo P, Dai Z, et al. clusterProfiler 4.0: A universal enrichment tool for interpreting omics data. Innovation (Camb). 2021;2(3):100141.

89. Cheng K, Brunius C, Fristedt R, Landberg R. An LC-QToF MS based method for untargeted metabolomics of human fecal samples. Metabolomics. 2020;16(4):46.

