## Supplementary material for "Butyrate ameliorates inflammation in colon biopsy samples of IBD patients and experimental colitis in mice involving RNA binding protein, AUF1-IL-27 axis and accelerating B1a to B10 polarization": Supplementary pdf.pdf

#### Materials and Methods

##### List of reagents used in the study

| | Cat# | Company |
|---|---|---|
| Monensin | M5273-5G | Sigma-Aldrich (Louis, MA, US) |
| Trypan blue | 15250-061 | Invitrogen (Waltham, MA, US) |
| Trizol | 9109 | Takara (Shiga, Japan) |
| Actinomycin D | A1410 | Sigma-Aldrich (Louis, MA, US) |
| Cyclohexamide | C4859 | Sigma-Aldrich (Louis, MA, US) |
| Cytofix/cytoperm | 51-2090KZ | BD Bioscience (NJ, USA) |
| Lipofectamine 2000 | 11668-027 | Invitrogen (Carlsbad, California) |
| Sheath fluid | 342003 | BD Bioscience (New Jersy, USA) |
| SyBr green | RR820A | Takara (Shiga, Japan) |
| cDNA kit | 6110A | Takara (Shiga, Japan) |
| KpnI RE<br>XhoI RE<br>HindIII | R3142<br>R0146<br>R0104 | NEB (Massachusetts, USA)<br>NEB (Massachusetts, USA)<br>NEB (Massachusetts, USA) |
| Ligase | M0202 | NEB (Massachusetts, USA) |
| Sodium butyrate | TC471-500G | Himedia (Kennett Square, USA) |
| Sodium acetate | GRM1012-500G | Himedia (Kennett Square, USA) |
| Sodium propionate | GRM6227-500G | Himedia (Kennett Square, USA) |
| Dextran sodium sulphate | 42867-25G | Sigma-Aldrich (Louis, MA, US) |
| paraformaldehyde | 30525-89-4 | Merck (New Jersy, USA) |
| Chloroform | DB3P730246 | Sigma-Aldrich (Louis, MA, US) |
| phenol | 105-95-2 | Merck (New Jersy, USA) |
| 2-propanol | 1.94524.0521 | Merck (New Jersy, USA) |
| Methanol | MB113-500ml | Himedia (Kennett Square, USA) |
| Ambion DNase1 | AM2222 | Invitrogen (Carlsbad, California) |

| | | |
|---|---|---|
| si-AUF1 | 12763S | Cell Signalling (Massachusetts, USA) |
| RPMI | 318000-022 | Thermofisher (Massachusetts, USA) |
| FBS | 2910154 | MP Biomedicals (California, USA) |
| Ficoll | 10771-500ml | Sigma-Aldrich (Louis, MA, US) |
| Sucrose | 61839805001730 | Merck (New Jersy, USA) |
| Hoechst 33342 | 62249 | Thermofisher (Massachusetts, USA) |
| Chow diet | (Harlan Teklad LM-485) | NIN (Hyderabad, India) |
| Diluent for DNA Extraction | MB228-500ML | Himedia (Kennett Square, USA) |
| PVDF membrane | 88518 | Invitrogen (Carlsbad, California) |
| Ripa Lysis buffer | 786-490 | GBiosciences (St. Louis, USA) |
| BCIP-NBT kit | 34042 | Thermofisher (Massachusetts, USA) |
| PBS | 823053 | MP Biomedicals (California, USA) |
| SPINeasy PCR Purification kit | 65380 | MP Biomedicals (California, USA) |
| Penicillin, streptomycin | 15140-122 | Thermofisher (Massachusetts, USA) |
| MTT | M2128 | Sigma-Aldrich (Louis, MA, US) |
| Fixable Viability Stain 780 | 565388 | BD Biosciences (NJ, USA) |
| TBS 10X | GCR-1T | GCC Biotech (New Delhi, India) |
| Tween 20 | RC1226 | GBiosciences (St. Louis, USA) |
| Tris HCL 6.8 | R178 | GBiosciences (St. Louis, USA) |
| Tris HCL 8.8 | R188 | GBiosciences (St. Louis, USA) |
| Running Buffer | ML042 | Himedia (Kennett Square, USA) |
| SuperSignal West Pico chemiluminescent substrate kit | 34580 | Thermofisher (Massachusetts, USA) |
| BCA Protein Assay Kit | 10741395 | Thermofisher (Massachusetts, USA) |
| Anti-mouse TNF-α ELISA kit | 558534 | BD Biosciences (NJ, USA) |
| Anti-mouse IL-10 ELISA kit | 555252 | BD Biosciences (NJ, USA) |
| Anti-mouse IL-6 ELISA kit | 555240 | BD Biosciences (NJ, USA) |

| | | |
|---|---|---|
| Anti-human TNF-α ELISA kit | 555157 | BD Biosciences (NJ, USA) |
| IL-27 ELISA | DY1834 | R&D Systems (Minnesota, USA) |

**List of antibodies used in the study**

| | Cat# | Company |
|---|---|---|
| Anti-AUF1 | 07-260 | Merck Millipore |
| Anti-IL-27 | PA5-115411 | Invitrogen (Carlsbad, California) |
| Anti-GAPDH | MA5-15738 | Invitrogen (Carlsbad, California) |
| Anti-β-actin | MA5-15739 | Invitrogen (Carlsbad, California) |
| Anti-CD3 | 100204 (clone 17A2) | BioLegend (SanDiego, California) |
| Anti-CD4 | 553052 (clone RM4-5) | BD Pharmingen (NJ, USA) |
| Anti-CD19 | 551001 (clone 1D3) | BD Pharmingen (NJ, USA) |
| Anti-CD5 | 553023 (clone 53-7.3) | BD Pharmingen (NJ, USA) |
| Anti-CD23 | 553139 (clone B3B4) | BD Pharmingen (NJ, USA) |
| Anti-CD1d | 123505 (clone 1B1) | BioLegend (SanDiego, California) |
| Anti-IL-10 | 554423 (clone SXC-1) | BD Pharmingen (NJ, USA) |
| Anti-CD16/CD32 (Fc Block) | 553142 (clone 2.4G2) | BD Pharmingen (NJ, USA) |
| APC streptavidin | 405207 | BioLegend (SanDiego, California) |
| FITC streptavidin | 405201 | BioLegend (SanDiego, California) |

All primers were purchased from IDT (Lowa, USA)

| | **Forward** | **Reverse** |
|---|---|---|
| AUF1 (mice) | 5'-AGAACGAGGAGGATGAAGGGA-3' | 5`-TGTGTCTGGAGAAAGGCCAC-3` |
| IL-27 p28 (mice) | 5'-ATGGACCCGGATCCTGAAGA-3' | 5'-TTATCCCTGGGCTCCCATCA-3' |
| EBI3 (mice) | 5'-CTCTCAAGTACCGACTCCGCTA-3' | 5'-CTGAGCTGACACCTGGATGCAA-3' |
| IL-10 (mice) | 5′-GGTTGCCAAGCCTTATCGGA-3′ | 5′-ACCTGCTCCACTGCCTTGCT-3′ |
| TNF-α(mice) | 5′-CATCTTCTCAAAATTCGAGTGACAA-3′ | 5′-TGGGAGTAGACAAGGTACAACCC-3′ |
| IFN-γ (mice) | 5′-TCAAGTGGCATAGATGTGGAAGAA-3′ | 5′-TGGCTCTGCAGGATTTTCATG-3′ |

| | | |
|---|---|---|
| GAPDH (mice) | 5`-AGAGAGGCCCAGCTACTCG-3` | 5`GGCACTGCACAAGAAGATGC-3 |
| AUF1 (human) | 5’-CGGCACAGCGGGAAGA-3’ | 5’-TGGTTCCAGTTTTGACTGGGG-3’ |
| IL-27 p28 (human) | 5’-GACCAAAGAGGCTGGGCCCC-3’ | 5’-TGGATGAGAGTGCTTTATTG-3’ |
| IL-10 (human) | 5’-GCCAAGCCTTGTCTGAGATGATCC-3’ | 5’-CTCACTCATGGCTTTGTAGATGCC-3’ |
| TNF-α (human) | 5’-GTGACAAGCCTGTAGCCCATGTTG-3’ | 5’-CTTGATGGCAGAGAGGAGGTTGAC-3’ |
| GAPDH (human) | 5`-GAGAAGGCTGGGGCTCATTT-3` | 5`-AGTGATGGCATGGACTGTGG-3` |
| IL-27 p28 (3’ UTR) | CCG CTCGAG TTCTAGACACCTAGCTTCAAGCCCTATGG | CGG GGTACC GGCCGGCCCGGGCTGGATGGCTTTATTA |

**Figure:**

#### Figure S1

**A**

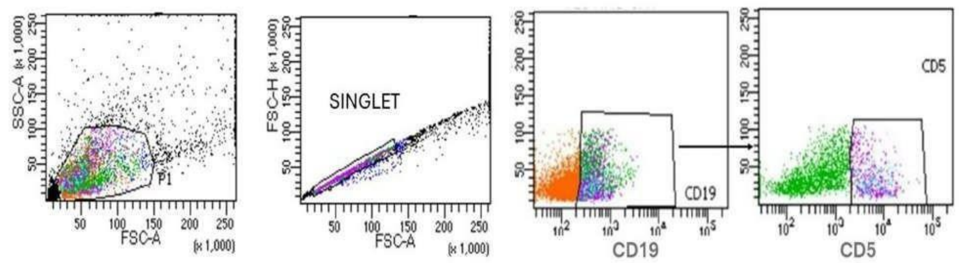


**B**

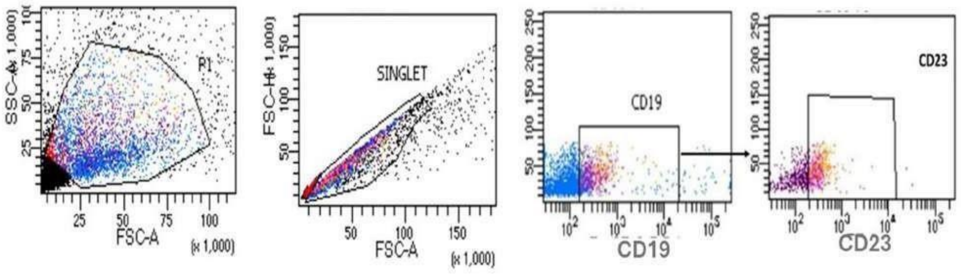


**Fig S1**: The splenocytes were treated with 100 µM of each SCFAs for 72 h and monensin was added to last 8 h of treatment. Thereafter cells were stained with antibodies against CD19, CD5 and CD23. Gating strategy of B1a cell (A) and B2 cell (B).

#### Figure S2

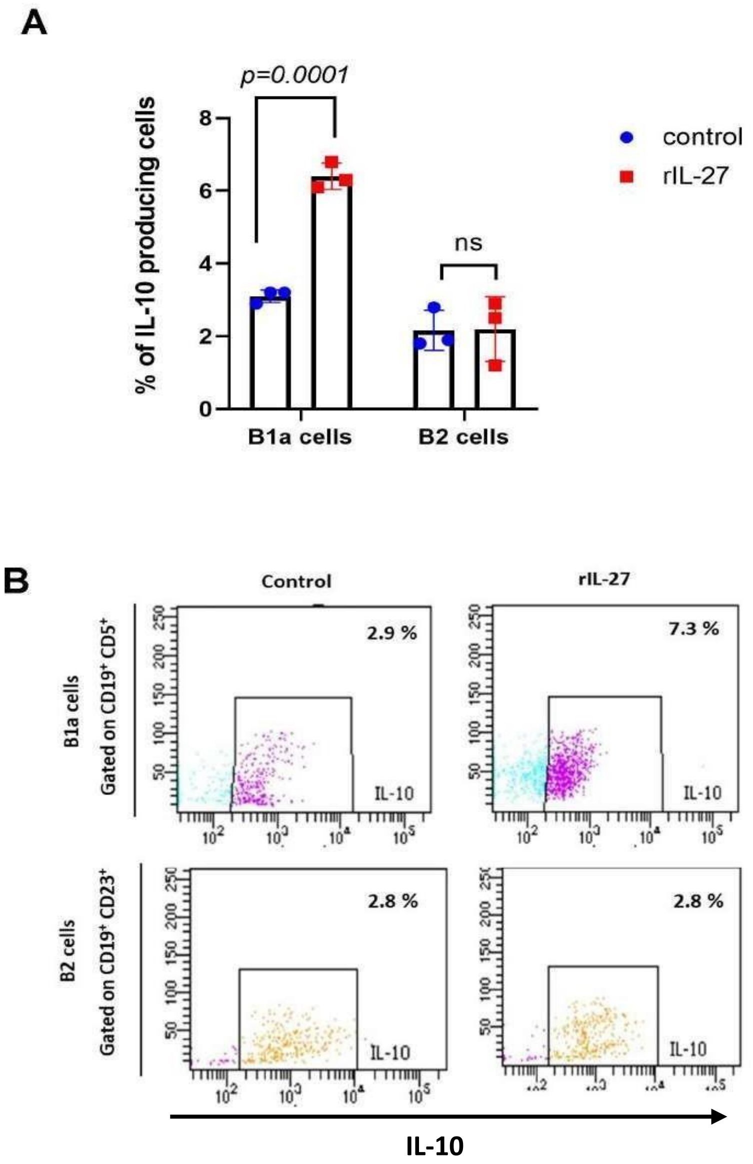


**Figure S2: Effect of rIL-27 on mouse splenocytes in terms of IL-10 production by Cd19$^{+}$CD5$^{+}$ (B1a), and Cd19$^{+}$ CD23$^{+}$ (B2) cells.**

Mouse splenocytes were treated with rIL-27 (20 ng/ml) for 72 h and frequencies of intracellular IL-10 producing Cd19$^{+}$CD5$^{+}$ **(**B1a) and Cd19$^{+}$ CD23$^{+}$ (B2) cells were assayed by flowcytometry (A). The representative dot plots are in (B). All experimented were repeated thrice (n=3) and represented as Mean±SE. *p < 0.05; **p < 0.01; ***p < 0.001 with respect to control calculated by student's t-test.

### Figure S3

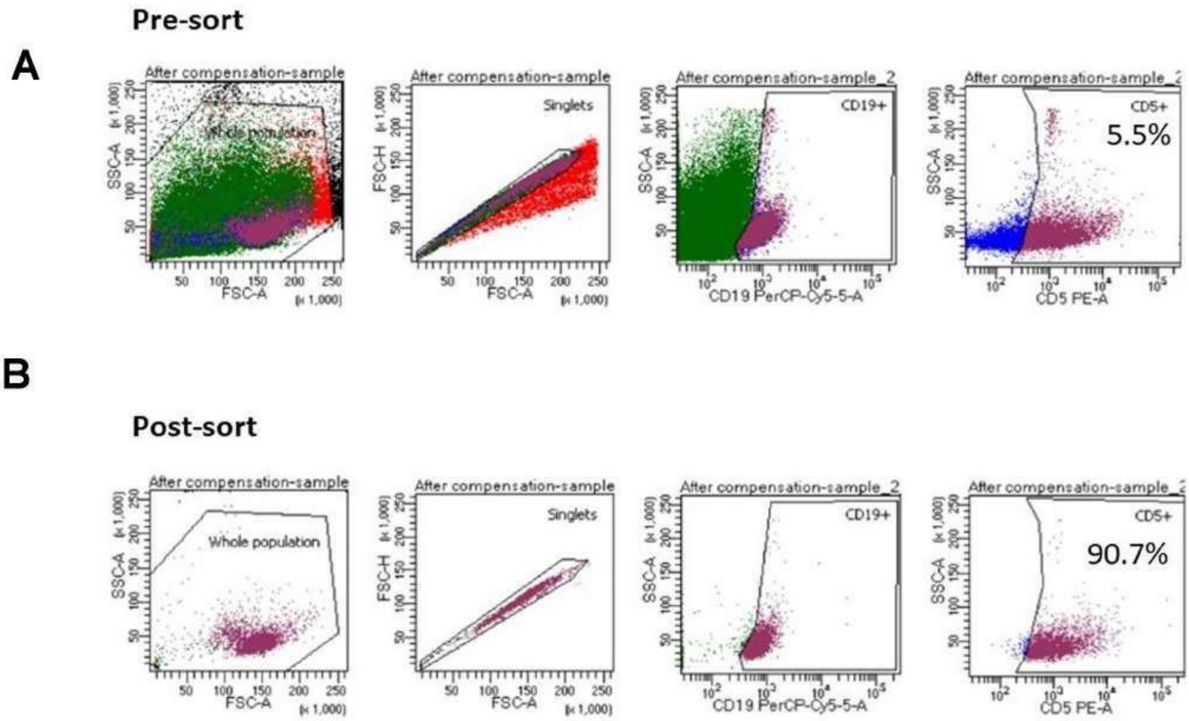


**Fig S3**: The splenocytes isolated from untreated mice and monensin was added followed by incubation for 8h. Thereafter cells were stained with antibodies against CD19 and CD5. The $CD19^{+}CD5^{+}$ cells were purified and were analyzed after sorting. Gating of B1a cell pre and post sort.

### Figure S4

A

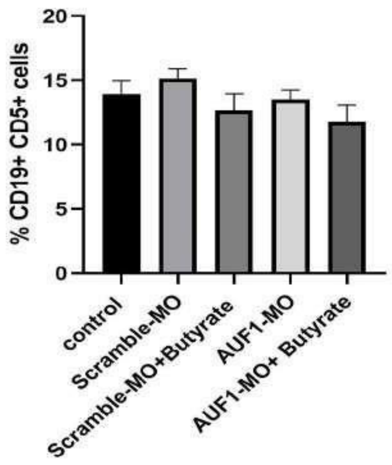


B

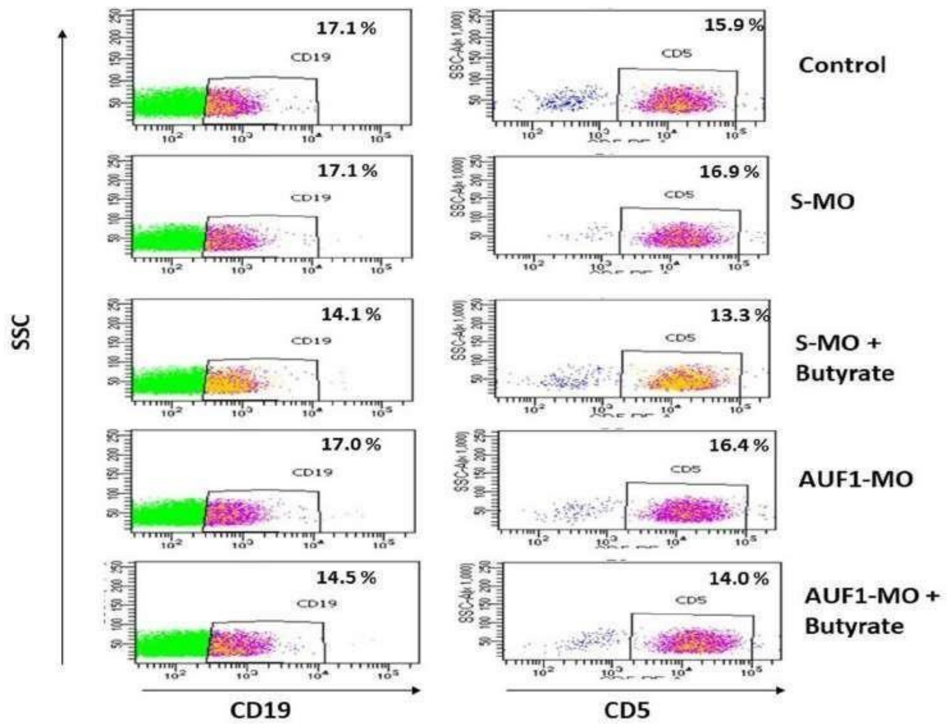


**Figure S4:** The splenocytes were isolated from control, S-MO, S-MO+butyrate, AUF1-MO and AUF1-MO+ butyrate mice, monensin was added and incubated for 8 h. Thereafter cells were stained with antibodies against CD19 and CD5. Percentage of $CD19^{+}CD5^{+}$ cells (A). Gating of $CD19^{+}CD5^{+}$ cells from splenocytes from control, S-MO, S-MO+butyrate, AUF1-MO and AUF1-MO+ butyrate mice (B).

### Figure S5

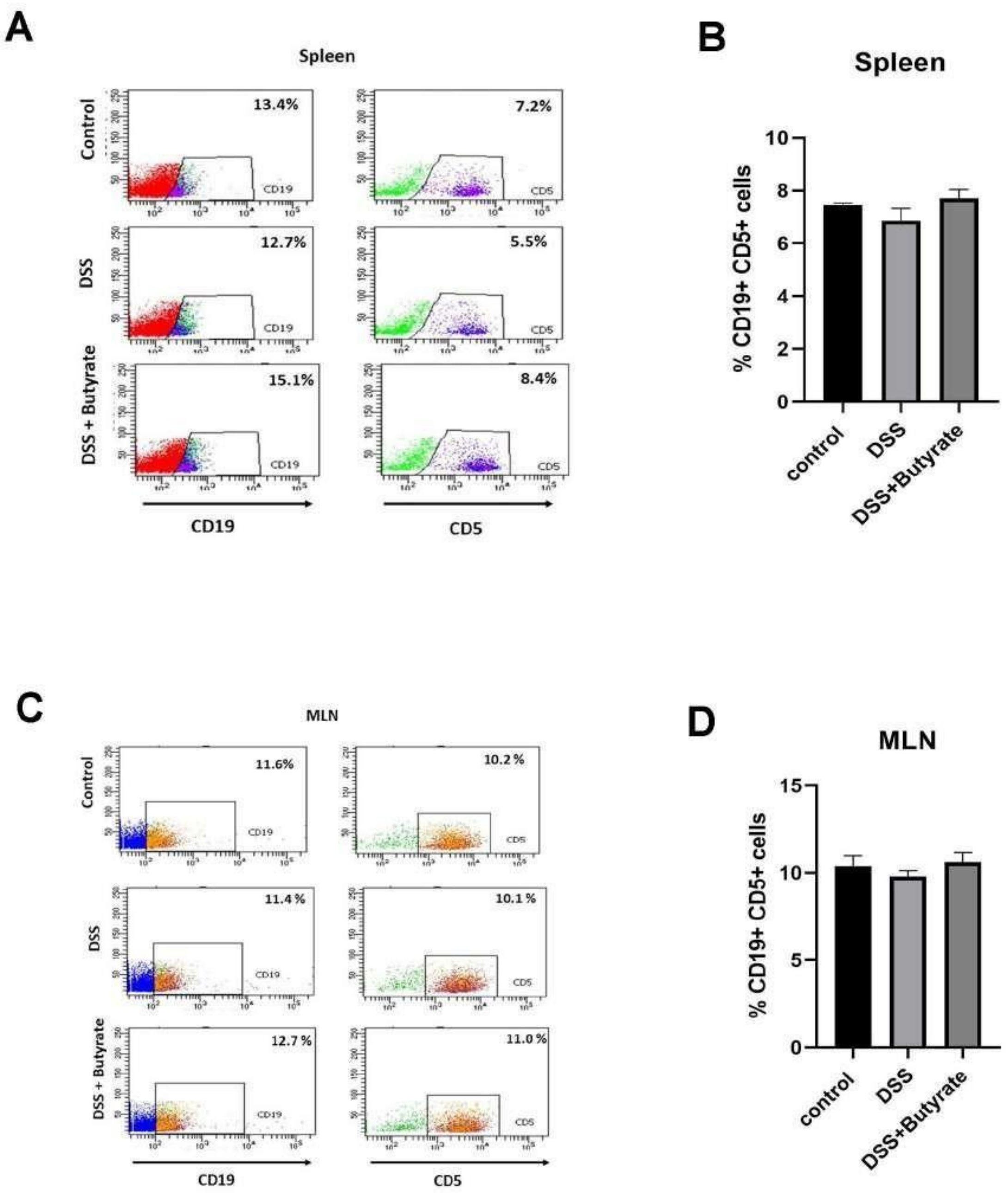


**Fig S5:** The splenocytes were isolated from control-mice, DSS-mice and DSS+Butyrate-mice, monensin was added and incubated for 8 h. Thereafter cells were stained with antibodies against CD19 and CD5. Gating of CD19$^{+}$CD5$^{+}$ cells (A) and percentage of CD19$^{+}$CD5$^{+}$ cells (B) in spleen of control-mice, DSS-mice and DSS + Butyrate-mice. Gating of CD19$^{+}$CD5$^{+}$ cells (C) and percentage of CD19$^{+}$CD5$^{+}$ cells (D) in MLN of control-mice, DSS-mice and DSS+Butyrate-mice.

#### Figure S6

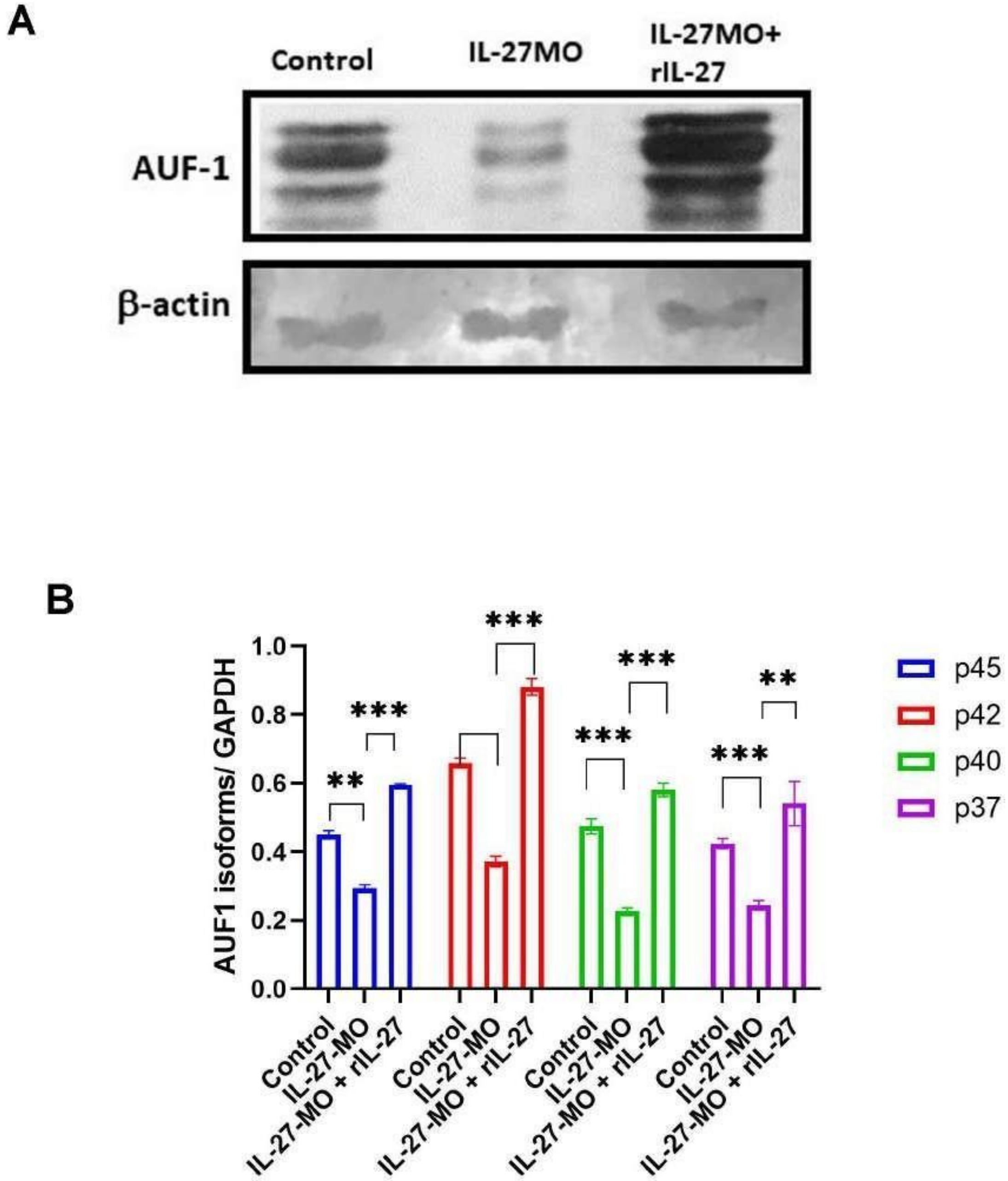


**Fig S6:** The splenocytes isolated from wild type mice were treated with either IL-27 MO or IL-27MO plus anti-IL-27mAb for 24h. There after the cells were lysed and western blot analysis was conducted using anti-AUF1 antibody and GAPDH antibody respectively (A). The densitometry analysis is represented (B).

### Figure S7

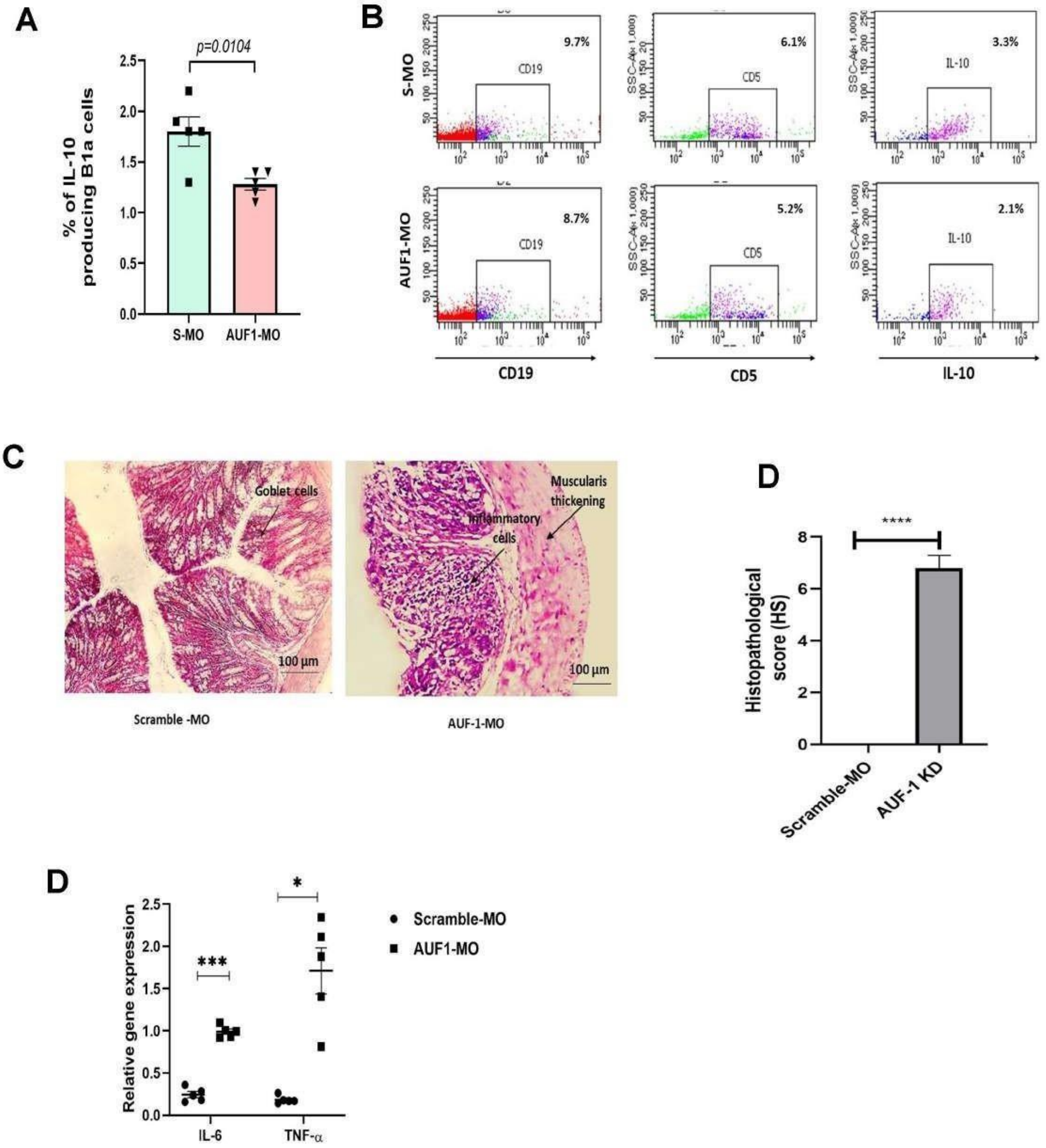


**Fig S7:** The cells from MLN were isolated from S-MO (wild type) and AUF1-MO (AUF1-KD) mice, monensin was added and incubated for 8 h. Thereafter cells were stained with antibodies against CD19, CD5 and IL-10. The percentage of $CD19^{+}CD5^{+}IL\text{-}10^{+}$ cells (A) and the gating strategy (B) in MLN of wild type and AUF1-KD mice are represented. The histology section (C) and histology score (D)of the colon tissue from wild type and AUF1-KD mice. The cytokines (TNF-α and IL-6) gene expression was determined from colon tissue of wild type and AUF1-KD mice (E).

#### References


1. Bora P, Gahurova L, Masek T, Hauserova A, Potesil D, Jansova D, et al. p38-MAPK-mediated translation regulation during early blastocyst development is required for primitive endoderm differentiation in mice. Commun Biol. 2021;4(1):788.
2. Panda AC, Martindale JL, Gorospe M. Polysome Fractionation to Analyze mRNA Distribution Profiles. Bio Protoc. 2017;7(3).
3. Abdelmohsen K, Tominaga-Yamanaka K, Srikantan S, Yoon JH, Kang MJ, Gorospe M. RNA-binding protein AUF1 represses Dicer expression. Nucleic Acids Res. 2012;40(22):11531-44.
4. Blake MS, Johnston KH, Russell-Jones GJ, Gotschlich EC. A rapid, sensitive method for detection of alkaline phosphatase-conjugated anti-antibody on Western blots. Anal Biochem. 1984;136(1):175-9.
5. Dobin A, Davis CA, Schlesinger F, Drenkow J, Zaleski C, Jha S, et al. STAR: ultrafast universal RNA-seq aligner. Bioinformatics. 2013;29(1):15-21.
6. Liao Y, Smyth GK, Shi W. featureCounts: an efficient general purpose program for assigning sequence reads to genomic features. Bioinformatics. 2014;30(7):923-30.
7. Love MI, Huber W, Anders S. Moderated estimation of fold change and dispersion for RNA-seq data with DESeq2. Genome Biol. 2014;15(12):550.
8. Wu T, Hu E, Xu S, Chen M, Guo P, Dai Z, et al. clusterProfiler 4.0: A universal enrichment tool for interpreting omics data. Innovation (Camb). 2021;2(3):100141.
9. Cheng K, Brunius C, Fristedt R, Landberg R. An LC-QToF MS based method for untargeted metabolomics of human fecal samples. Metabolomics. 2020;16(4):46.
